# Inactivation of PRMT1 inhibits cardiac fibrosis via transcriptional regulation and perturbation of FBL activity in fibroblast-to-myofibroblast transition

**DOI:** 10.1101/2024.05.24.595777

**Authors:** Jiang Qian, Olan Jackson-Weaver, Yudao Shen, Jian Jin, Yibu Chen, Meng Li, Ram Kumar Subramanyan, Jian Xu

## Abstract

Cardiac fibrosis is a recognized cause of morbidity and mortality, yet effective pharmacological therapy that directly targets the fibrotic process remains lacking. Here we surveyed a group of methyltransferases known as protein arginine methyltransferases (PRMT) and demonstrated that PRMT1, which is the most highly expressed PRMT in the heart, was upregulated in activated cardiac fibroblasts, or myofibroblasts, in failing hearts. Deleting Prmt1 specifically in myofibroblasts or treating systemically with the PRMT1 inhibitor MS023 blocked myofibroblast formation, leading to a significant reduction in cardiac fibrosis and improvement in cardiac function in both acute and chronic heart injury models that manifest pervasive cardiac fibrosis. PRMT1 promoted the transition of cardiac fibroblasts to myofibroblasts by regulating transcription and epigenetic status. Additionally, PRMT1 methylated a key nucleolar protein fibrillarin 1 (FBL) and regulated nucleoli morphology and function during fibroblast fate transition. We further demonstrated a previously unrecognized requirement for FBL in myofibroblasts formation, by regulating myofibroblast gene induction and contractile force generation.

## INTRODUCTION

Cardiac fibrosis, characterized by excessive deposition of extracellular matrix, increases myocardial stiffness, disrupts cardiac conduction, and impairs systolic and diastolic function^1–3^. It is a significant contributor to heart failure and increases morbidity and mortality of other cardiovascular conditions. Unfortunately, there is no approved therapies that could specifically target the fibrotic process.

Cardiac fibroblasts are the major matrix-producing cells within the heart. In pro-fibrotic conditions like hypertensive heart disease, cardiac fibroblasts become activated and transition into myofibroblasts. These myofibroblasts express highly contractile proteins, deposit matrix components, and secrete pro-inflammatory cytokines^1, 2^, ultimately leading to cardiac fibrosis and dysfunction. The transition of cardiac fibroblasts to myofibroblasts can be triggered by extracellular stimuli such as growth factors and cytokines, communication with other cardiac cell types^4, 5^, or intrinsic mechanisms within fibroblasts themselves, involving signaling, transcriptional and translational processes^2^.

Recently, therapeutic strategies based on transcriptional modulators and epigenetic enzymes have been explored to alleviate fibrosis in pre-clinical trials of cardiac, pulmonary, hepatic, and renal diseases^3, 6^. Here we surveyed a group of methyltransferases known as protein arginine methyltransferases (PRMT), a family of nine enzymes responsible for methylation of arginine residues in proteins including histones, transcription factors and signaling mediators.

Specifically, we focused on PRMT1, which is the most abundant PRMT in the heart. PRMT1 expression was elevated in activated cardiac fibroblasts, or myofibroblasts, in failing hearts. Deleting Prmt1 specifically in myofibroblasts or treating systemically with the PRMT1 inhibitor MS023 blocked myofibroblast formation, leading to a significant reduction in cardiac fibrosis and improvement in cardiac function in both acute and chronic heart injury models that manifest extensive cardiac fibrosis. PRMT1 was found to promote the transition of fibroblasts to myofibroblasts by regulating transcription and epigenetic status. Additionally, PRMT1 methylated a key structural constituent of nucleoli, fibrillarin 1 (FBL), regulating nucleoli morphology and function to alter protein synthesis during fibroblast fate transition to myofibroblast. We further demonstrated a previously unrecognized role for FBL in cardiac fibroblasts fate transition, by regulating myofibroblast gene induction and contractile force generation.

## RESULTS

### PRMT1 expression increased in the interstitial fibroblasts of failing hearts

Cardiac remodeling, driven by excessive pressure and volume load in conditions such as hypertension, initiates a detrimental cycle of progressive cardiac dysfunction culminating in heart failure. In a previous study by Yang et al., the coding transcriptome in the left ventricle of failing human hearts from ischemic and non-ischemic diseases was quantitatively analyzed, revealing significant differences in mRNA expression patterns^7^. We conducted secondary analysis of transcriptomic data from these clinical cohorts and assess the expression profile of protein arginine methyltransferase (PRMT) family of enzymes. PRMT1 is the highest expressing PRMTs in the heart (Fig. 1A). The expression of PRMT1 mRNA increased in heart failure patients with ischemic and non-ischemic conditions (Fig. 1A), suggesting regulatory roles of PRMT1 in the context of cardiac reverse remodeling.

**Figure 1.**
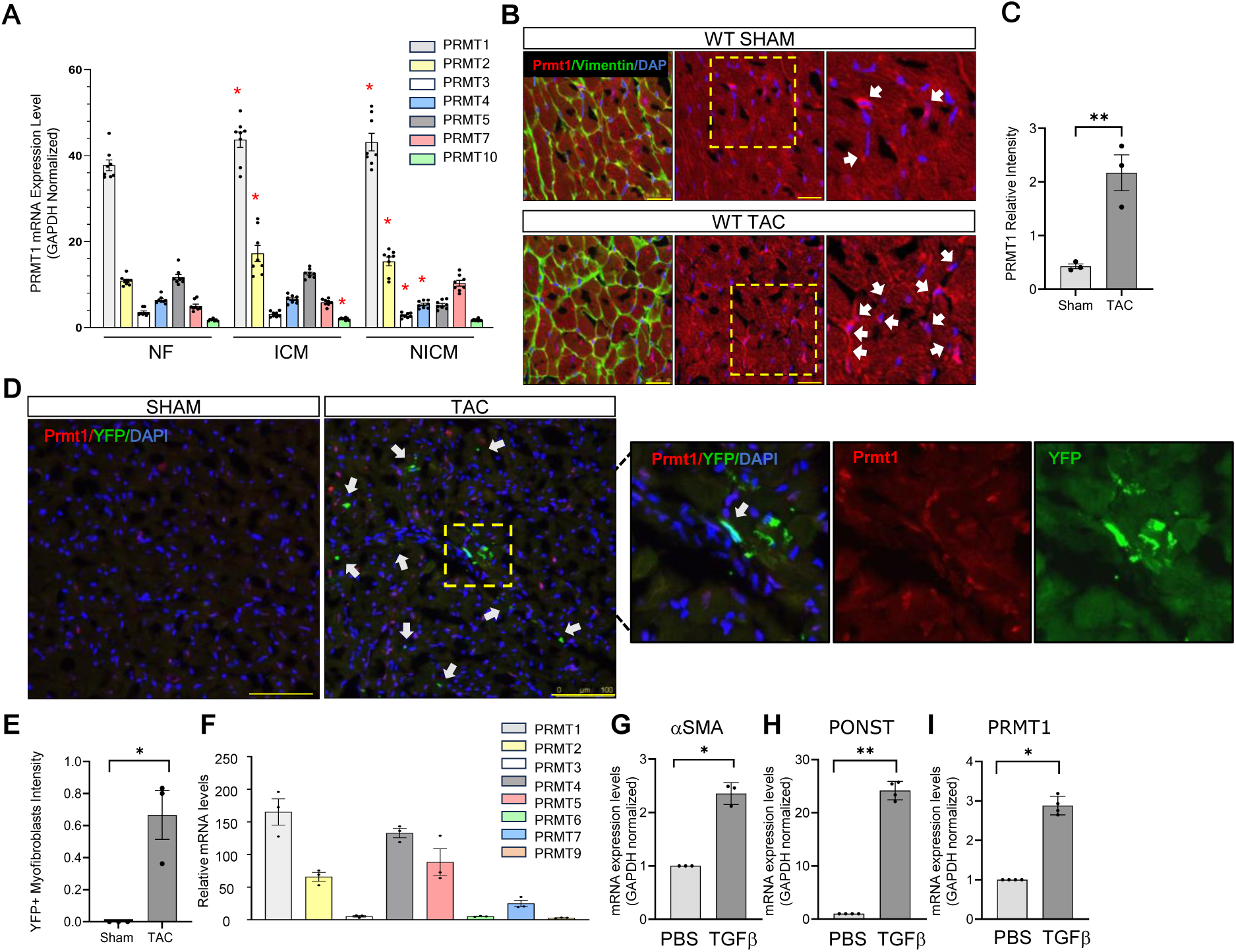
PRMT1 expression positively correlates with disease progression in heart failure. **A**. PRMT1 expression increased in heart failure patients of ischemic cardiomyopathy (ICM) or non-ischemic cardiomyopathy (NICM), comparing with non-failing hearts (NF), through secondary analysis of a published clinical cohort. **B-C.** PRMT1 expression (red) increased in wild type mouse heart at 8 weeks post-TAC surgery when compared to sham-operated controls. PRMT1 colocalized with vimentin (green), which labels cardiac fibroblasts. The level of increase in PRMT1 protein expression was quantified and shown in C. **D-E.** Elevated PRMT1 expression (red) was identified in cardiac myofibroblasts in TAC-induced heart failure mouse model. Cardiac myofibroblasts were labeled by YFP expression (green) using a genetic lineage tracing model, *Pn^MCM^;ROSA^STOP-YFP^*. The increase in the number of YFP+ myofibroblasts in TAC vs. Sham group was quantified and shown in E. **F.** The mRNA expression of PRMTs in human cardiac fibroblasts. **G-I.** PRMT1 mRNA expression increased in human cardiac fibroblasts stimulated with TGFβ to undergo fibroblast to myofibroblast fate transition. The mRNA levels of PRMT1 and myofibroblast marker genes *aSMA* and *POSTN* was assessed with RT-PCR. Data are shown as mean ± SEM and representative of three biological replicates. All reactions were performed in technical triplicates and normalized to GAPDH. *: P<0.05, **: P<0.01. Scale bar=100μm.

We then used a mouse model of chronic heart failure induced by transverse aortic constriction (TAC) surgery. In these mice, PRMT1 expression was elevated in the failing hearts compared to sham-operated non-failing hearts (Fig. 1B, 1C). The expression of PRMT1 revealed by immunohistostaining was enriched in the interstitium, where cardiac fibroblasts represent the major cell population. Given the significant contribution of fibroblast activation to fibrosis during heart failure pathogenesis, we specifically identified activated fibroblasts, or myofibroblasts, in post-TAC failing mouse hearts using a mouse model that genetically labels myofibroblasts with a yellow fluorescent protein (YFP) reporter to facilitate their identification (Fig. 1D). To this end, the myofibroblast-specific tamoxifen-inducible Cre line *Postn^MCM^* ^8, 9^ was bred with the YFP reporter line, *Rosa^STOP-YFP^*, enabling us to identify and trace activated fibroblast cells by YFP expression (Supplemental Fig. S1). These YFP-positive myofibroblasts exhibited heightened expression of PRMT1 (Fig. 1D, Fig. 1E).

To further investigate the relationship between PRMT1 and fibroblast activation, we used primary human cardiac fibroblasts (hCF). Similarly to human heart, PRMT1 is the most abundant PRMT in human cardiac fibroblasts (Fig. 1F). We stimulated hCF with TGFβ, a potent inducer of fibroblast activation that leads to fate transition into myofibroblasts^4, 9, 10^. After 48 hours of TGFβ stimulation, hCFs underwent transition into myofibroblasts, as indicated by elevated expression of myofibroblast marker genes, *αSMA* and *POSTN* (Fig. 1G-H). Concurrently, PRMT1 expression significantly increased (Fig. 1I), further supporting the notion that activated fibroblasts express higher levels of PRMT1.

### Deletion of PRMT1 specifically in activated fibroblasts prevented their formation and reduced cardiac fibrosis

To assess the functional importance of PRMT1 in activated fibroblasts in vivo, we employed mouse genetic models to delete *Prmt1* in activated fibroblasts, or myofibroblasts, by crossing the myofibroblast-specific tamoxifen-inducible Cre line, *Postn^MCM^*, with *Prmt1^fl/fl^* and *Rosa^STOP-YFP^* mice (Fig. 2A)^8^. The *Rosa^STOP-YFP^* reporter enabled us to identify activated fibroblast cells through YFP expression (Supplement Fig. S2), since myofibroblast markers such as αSMA, periostin and vimentin are all expressed by multiple cell types. To induce cardiac fibrosis, we administered isoproterenol (ISO), a nonselective beta-adrenergic agonist, via osmotic mini-pump for two weeks (Fig. 2A). PRMT1 deletion was demonstrated by immunostaining showing the loss of PRMT1 protein in YFP+ myofibroblasts from ISO-treated CKO hearts (Fig. 2B, 2C, white arrows). Following ISO administration, we observed a significant increase in myofibroblasts, the main drivers of cardiac fibrosis, indicated by increased number of YFP^+^ cells in ISO-treated group, when compared to the saline-treated groups, which showed only a few myofibroblasts (Fig. 2D). Upon *Prmt1* deletion in activated fibroblasts, ISO-stimulated formation of myofibroblasts was inhibited (Fig. 2D, 2E). Consequently, PRMT1 deletion inhibited the development of cardiac fibrosis in the ISO-treated groups (Fig. 2F, 2G). Functionally, ISO administration resulted in apparent cardiac hypertrophy, indicated by an increase in heart weight to body weight ratio, and lung congestion from diastolic dysfunction, indicated by an increased in lung weight to body weight ratio (Fig. 2H, 2I). However, these effects were attenuated when *Prmt1* was deleted in activated fibroblasts, suggesting the beneficial effects of PRMT1 deletion in ameliorating cardiac fibrosis and functional decline (Fig. 2H, 2I, black vs. grey bars). These findings demonstrate the critical role of PRMT1 in cardiac fibroblasts for transition into myofibroblasts and the development of cardiac fibrosis in vivo.

**Figure 2.**
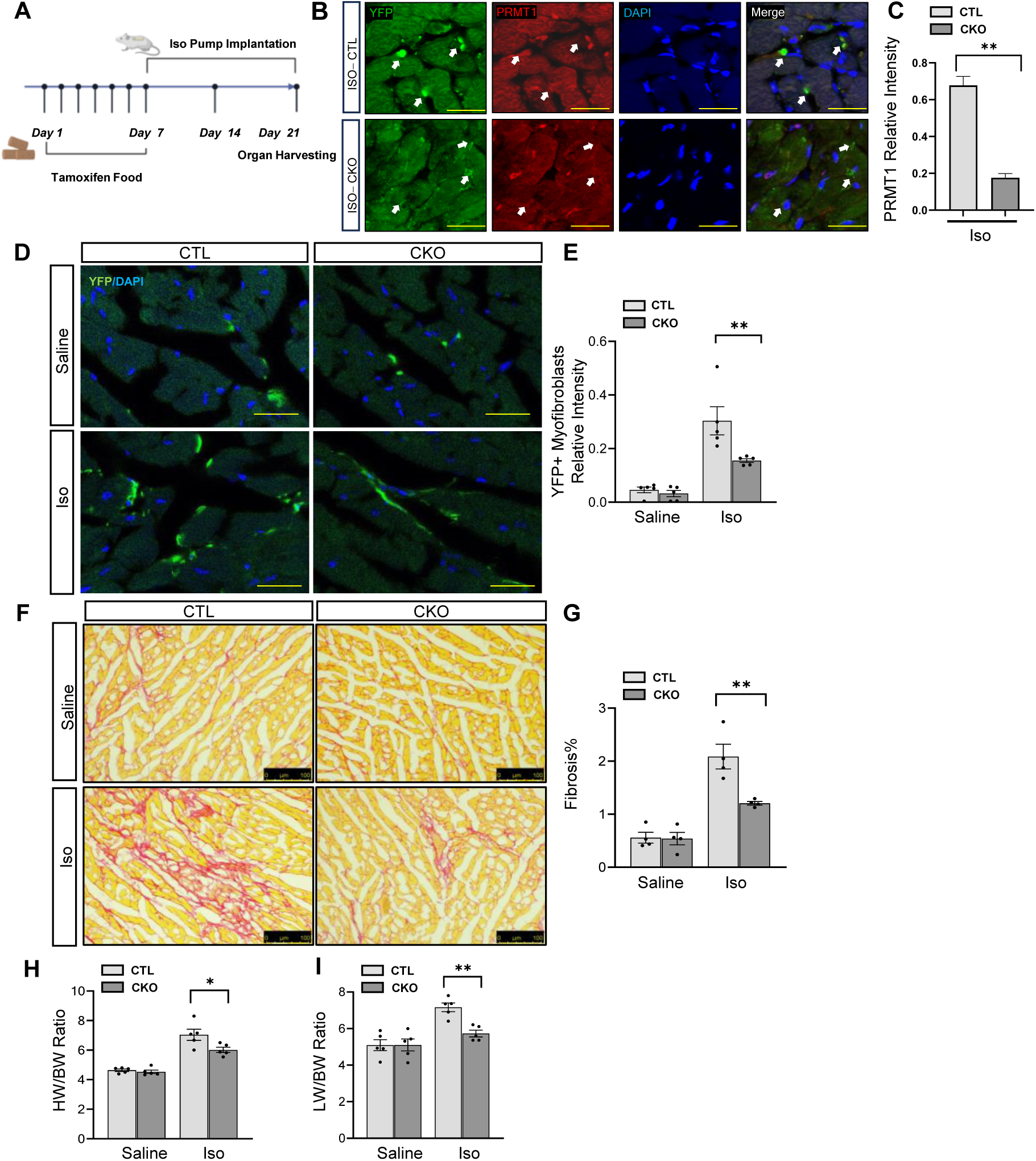
Myofibroblast-specific deletion of PRMT1 blocked myofibroblast formation and decreased cardiac fibrosis. **A.** Experimental timeline of isoproterenol (ISO)-induced mouse cardiac fibrosis by osmotic minipump infusion. **B-C.** PRMT1 protein expression was ablated in myofibroblasts of *Prmt1* knockout mouse heart. Myofibroblasts in the control (CTL, *Pn^MCM^;ROSA^STOP-YFP^*) and conditional *Prmt1* knockout (CKO, *Pn^MCM^;ROSA^STOP-YFP^;Prmt1^fl/fl^*) mice receiving ISO treatment were revealed by YFP expression with immunohistostaining. PRMT1 signal was lost in YFP+ cells from CKO hearts, in contrast to YPF+ cells from CTL hearts. PRMT1 signal in other cardiac cell types remained intact. PRMT1 signal in YPF+ myofibroblasts was quantified by Image J software and shown in C. **D-E.** The increase in cardiac myofibroblasts was inhibited by *Prmt1* deletion. Myofibroblasts were revealed by YFP expression using immunohistostaining of mouse heart tissue. Quantification of YFP+ myofibroblasts was conducted by Image J and shown in E. **F-G.** ISO-induced cardiac fibrosis was inhibited in CKO mice, indicated by Picrosirius red staining. Quantification of Picrosirius red signal was conducted by Image J and shown in G. **H.** ISO-induced increase of heart weight / body weight (HW/BW) ratio was attenuated in CKO mice. **I.** ISO-induced increase of lung weight / body weight (LW/BW) ratio was inhibited in CKO mice. *: P<0.05, **: P<0.01. Scale bar 100μm.

### Fibroblast-specific deletion of PRMT1 decreased cardiac fibrosis in TAC-induced heart failure model

To investigate the potential protective effects of PRMT1 deletion in heart failure, we utilized TAC surgery induced chronic heart failure model (Fig. 3A)^11^. The TAC procedure leads to sustained pressure overload that progresses into heart failure over a period of 3 months. TAC surgeries were performed on two groups of mice: *Postn^MCM^: Rosa^STOP-YFP^* (control, CTL) and *Postn^MCM^: Prmt1^fl/fl^ : Rosa^STOP-YFP^* (*Prmt1* CKO) (Fig. 3A). We confirmed the deletion of *Prmt1* by loss of PRMT1 protein expression in YPF+ myofibroblasts from the CKO heart (Fig. 3B, 3C). In the control mice, TAC-induced chronic injury resulted in a significant increase of YFP+ myofibroblasts (Fig. 3D, 3E), leading to cardiac fibrosis within the injured myocardium (Fig. 3F. 3G). Remarkably, the total myofibroblast counts in post-TAC *Prmt1* CKO hearts were significantly reduced (Fig. 3D, 3E), along with a dramatic decrease of cardiac fibrosis (Fig. 3F. 3G), when compared with control hearts after TAC. Furthermore, TAC-induced pressure overload leads to hypertrophic growth, a primary indicator of maladaptive heart remodeling^11^. This hypertrophic growth was notably attenuated when *Prmt1* was deleted, suggesting that the reduction in myofibroblast formation in *Prmt1* CKO also attenuated maladaptive remodeling in the cardiomyocytes (Fig. 3H).

**Figure 3.**
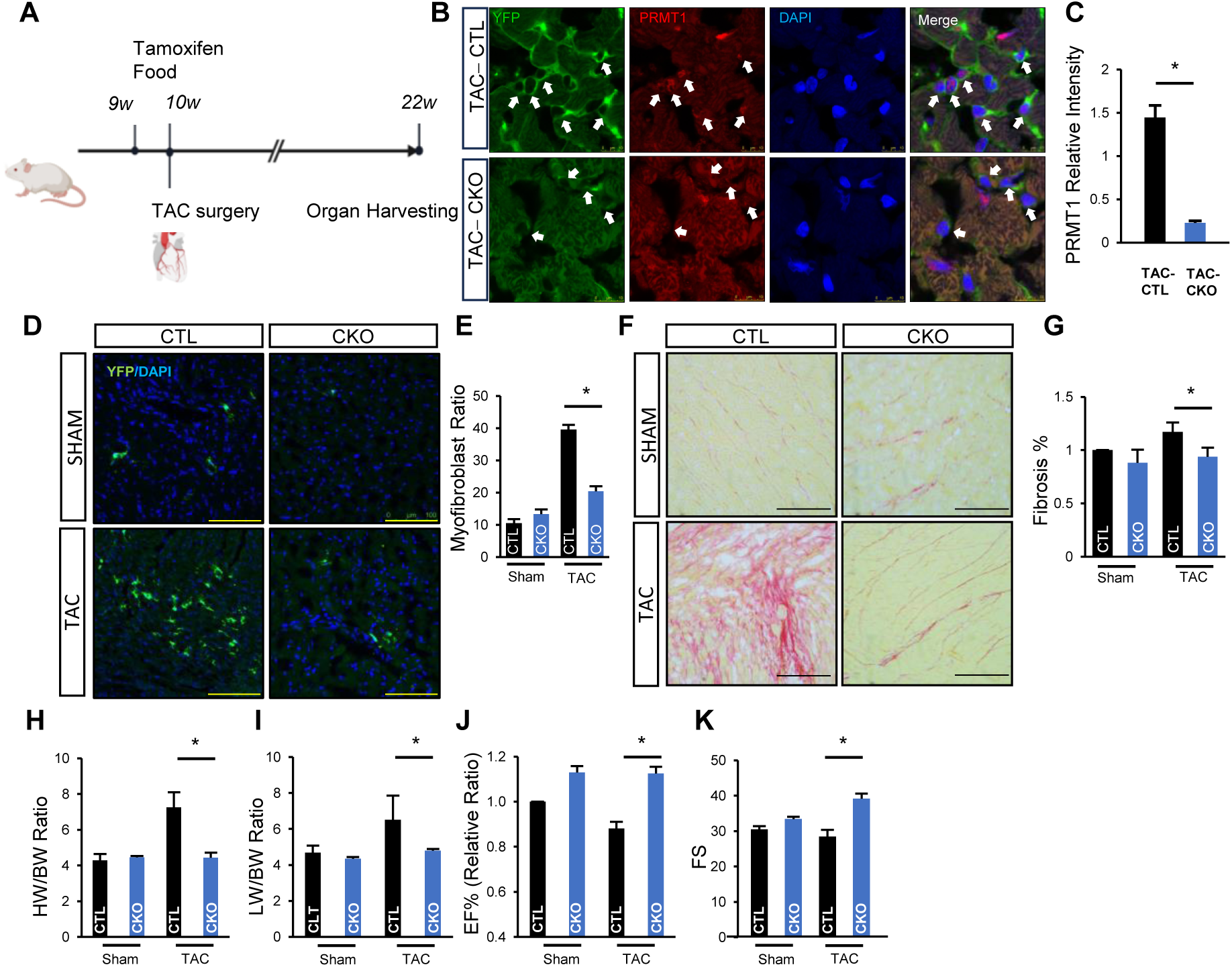
Inactivation of PRMT1 decreased cardiac fibrosis in TAC-induced heart failure. **A.** Experimental timeline of TAC surgery-induced chronic heart failure. **B-C.** PRMT1 protein expression was abolished in myofibroblasts of *Prmt1* CKO mouse heart. Myofibroblasts in the control (CTL) and *Prmt1* knockout (CKO) mice following TAC surgery were revealed by YFP expression with immunohistostaining. PRMT1 signal was lost in YFP+ cells from CKO hearts, in contrast to YPF+ cells from CTL hearts. PRMT1 signal in other cardiac cell types remained intact. PRMT1 signal in YPF+ myofibroblasts was quantified by Image J and shown in C. **D-E.** The increased cardiac myofibroblast formation as a result of TAC-induced heart failure was inhibited by *Prmt1* deletion. Myofibroblasts in mouse heart tissue were revealed by YFP expression using immunohistostaining. Quantification of YFP+ myofibroblasts was conducted by Image J and shown in E. **F-G.** TAC-induced cardiac fibrosis was repressed in CKO mice, indicated by reduction in Picrosirius red staining. Quantification of Picrosirius red signal was conducted by Image J and shown in G. **H-I.** TAC-induced increase in HW/BW ratio (H) and LW/BW ratio (I) was attenuated in CKO mice. **J-K.** Contractile functional decline was ameliorated in CKO mice at 12 weeks after TAC, indicated by echocardiographic assessment of ejection fractioning (EF%) in J and fractional shortening (FS) in K. Results are reported as the mean ± SEM from 6 mice in each group. *P < 0.05 using t-test. Scale bar=100μm.

To assess the decline in heart function following TAC-induced pressure overload, we analyzed lung congestion, as an indicator of diastolic dysfunction, and monitored cardiac function monthly for up to 12 weeks post-surgery. Similar to the isoproterenol-induced heart injury model, TAC surgery induced significant lung congestion in control mice, evidenced by lung weight measurements, but not in *Prmt1* CKO group (Fig. 3I). Ejection fraction (EF) and fraction shortening (FS), indexes of left ventricular systolic function, demonstrated significant decline in control mice but remain preserved in *Prmt1* CKO mice after TAC surgery (Fig. 3J, 3K). Collectively, our findings demonstrate that deletion of PRMT1 in activated fibroblasts led to a reduced myofibroblast formation and diminished cardiac fibrosis, along with preserved cardiac function, indicating PRMT1 as a potential target in preventing fibrosis and heart failure.

### Inhibition of PRMT1 activity using MS023 alleviates cardiac fibrosis in TAC-induced heart failure

PRMT1 is a druggable target, and one of its well-characterized inhibitors, MS023, effectively inhibits PRMT1 activity ex vivo at the nanogram level^12^. To investigate whether MS023 could provide protective effects in pressure overload-induced heart failure, we induced heart failure in wildtype C57B6 mice with TAC surgery and confirmed functional decline at 8 weeks after surgery when EF% significantly declined from the baseline (Supplement Fig. S3). Then, we administered MS023 (30 mg/kg/day) or saline as control daily for 4 weeks via intraperitoneal injection and monitored heart function in these mice by echocardiography (Fig 4A). MS023 treated group presented a significant reduction in cardiac fibrosis when compared to saline-treated group (Fig. 4B-C). MS023 treatment also started to improve cardiac function significantly at week 3 and sustained at week 4, while cardiac function in the saline treated group continued to decline (Fig. 4D-E).

**Figure 4.**
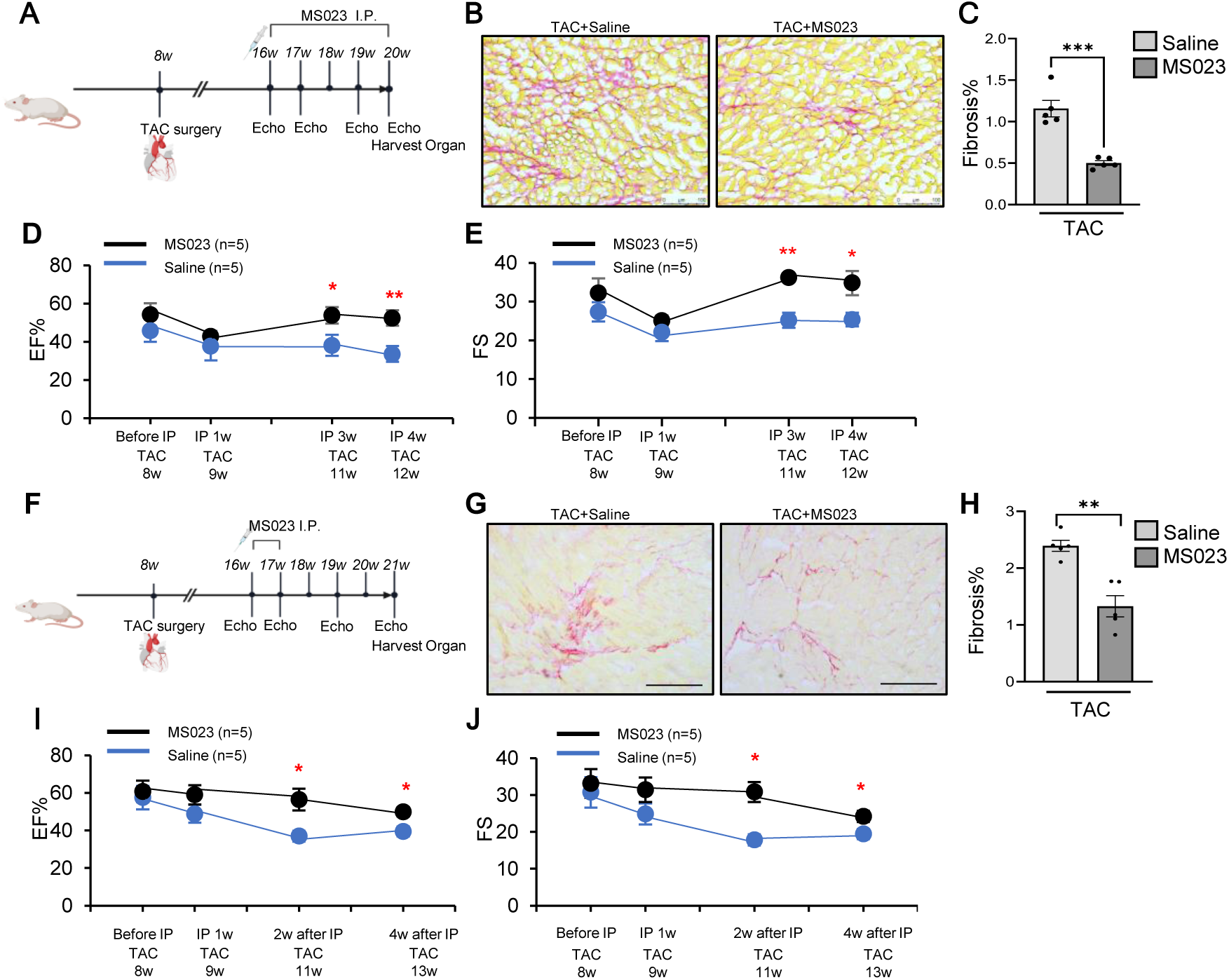
MS023 prevented cardiac fibrosis and dysfunction in TAC-induced heart failure. **A.** Experimental timeline using low dosage of MS023 (30μg/kg/day) treatment for 4 weeks in wild type mice that manifested cardiac dysfunction following TAC surgery (n=8 for MS023, n=5 for saline). **B-C.** TAC-induced cardiac fibrosis was reduced by MS023 treatment, indicated by decreased Picrosirius red staining. Quantification of Picrosirius red signal was conducted by Image J and shown in C. **D-E.** Contractile functional was improved by MS023 treatment at week 3 and 4, indicated by echocardiographic assessment of ejection fractioning (EF%) in B and fractional shortening (FS) in C. **F.** Experimental timeline using high dosage of MS023 (100μg/kg/day) treatment for 1 week in wild type mice that manifested cardiac dysfunction following TAC surgery (n=8 for MS023, n=5 for saline). **G-H.** TAC-induced cardiac fibrosis was reduced by MS023 treatment, indicated by decreased Picrosirius red staining. Quantification of Picrosirius red signal was conducted by Image J and shown in G. **I-J.** Contractile functional decline was alleviated by MS023 treatment and maintained for 4 weeks after treatment, indicated by echocardiographic assessment of ejection fractioning (EF%) in E and fractional shortening (FS) in F. Scale bar=100μm. *P < 0.05 using t-test.

Additionally, we tested a high-dose treatment regimen using daily administration of MS023 (100 mg/kg/day) for 7 days via intraperitoneal injection. Cardiac function was assessed by echocardiography before MS023 treatment and monitored biweekly after the treatment for 4 weeks (Fig 4F). MS023 treated group demonstrated a significant reduction of fibrosis when compared to saline-treated group (Fig. 4G-H). Remarkably, MS023 improved cardiac contractile function when compared to saline-treated group, and the beneficial effects were sustained for 4 weeks (Fig. 4I-J). To assess the safety of MS023 treatment in mice, we further performed histological analysis of MS023-treated mice and did not observe pathological changes in major organs (Supplement Fig. S4).

Taken together, MS023 treatment reduced fibrosis and improved cardiac function in TAC-induced chronic heart failure, suggesting that pharmacological inhibition of PRMT1 demonstrates protective effects, similar as genetic deletion of *Prmt1* in cardiac fibroblasts.

### PRMT1 is essential for the transition of cardiac fibroblast to myofibroblast

To investigate the mechanistic role of PRMT1 in the transition of cardiac fibroblasts to myofibroblasts, we isolated fibroblasts from adult mouse hearts and transfected them with siRNAs targeting PRMT1 or scrambled control siRNA. The mouse cardiac fibroblasts (mCF) were then treated with TGFβ to stimulate fibroblast activation and fate transition. TGFβ treatment induced a substantial increase in the expression of the myofibroblast marker αSMA, accompanied by characteristic stress fiber formation in control siRNA-transfected mCF, but not in PRMT1 siRNA-transfected group (Fig. 5A-B). PRMT1 was induced by TGFβ, similar as in hCF (Fig. 5C, 1I). Moreover, increased expression of myofibroblast signature genes, including *Col1α1, Col1α2, Sparc, Fn EDA, Fn1, Postn,* and *αSMA* was observed in control siRNA-transfected fibroblasts after exposure to TGFβ, but not in PRMT1 siRNA-transfected fibroblasts with the same TGFβ treatment (Fig. 5C-J).

**Figure 5.**
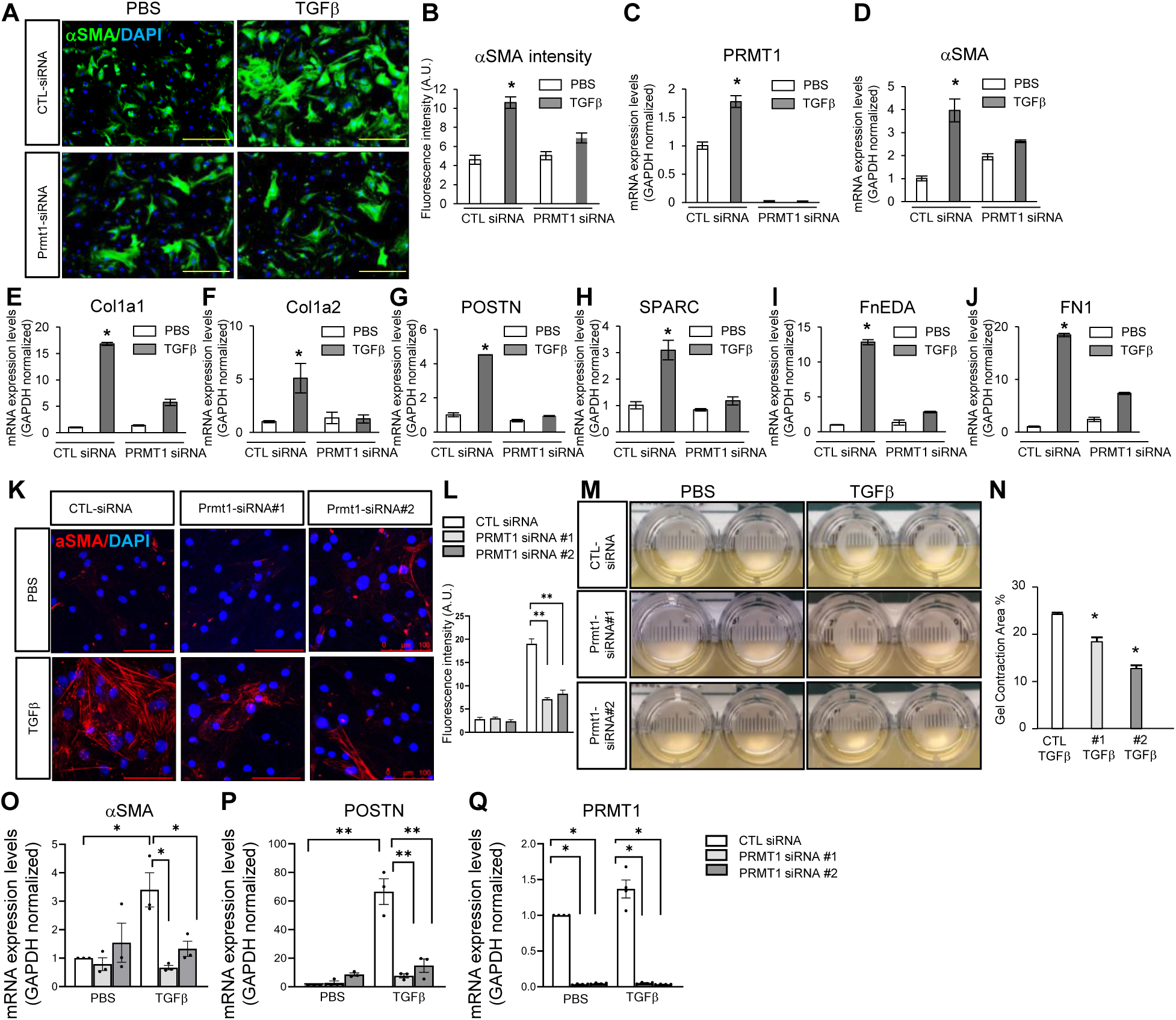
Depletion of PRMT1 inhibited cardiac fibroblasts to myofibroblasts fate transition. **A-B.** PRMT1 depletion suppressed TGFβ-induced expression of myofibroblast marker αSMA in mouse cardiac fibroblasts (mCF). Immunofluorescent signal of myofibroblast marker αSMA in cultured mCF with TGFβ or PBS treatment after PRMT1 siRNA or control (CTL) siRNA transfection was quantified and shown in B. **C-J.** The induction of myofibroblast marker genes *aSMA, Col1a1, Col1a2, Postn, Sparc, FnEDA* and *Fn* was repressed by PRMT1 knockdown in mCFs stimulated with TGFβ to undergo cardiac fibroblast to myofibroblast fate transition. The mRNA levels of PRMT1 and myofibroblast marker genes were assessed by RT-PCR. Data are shown as mean ± SEM and representative of three biological replicates. All reactions were performed in technical triplicates and normalized to GAPDH. **K-L.** In human cardiac fibroblasts (hCF), PRMT1 depletion suppressed TGFβ-induced expression of myofibroblast marker αSMA. Immunofluorescent signal of αSMA in cultured hCF transfected with two independent PRMT1 siRNA (#1 and #2) or control (CTL) siRNA followed by TGFβ or PBS treatment was quantified and shown in L. **M-N.** PRMT1 depletion inhibited contractile force generation. Collagen contractility assay of cultured hCF treated with TGFβ to induce cardiac fibroblast to myofibroblast fate transition or PBS as control. The area of gel contraction was quantified and shown in N. **O-Q.** The induction of myofibroblast marker genes *aSMA* and *Postn* was repressed by PRMT1 knockdown in hCFs stimulated with TGFβ. The mRNA levels of *Prmt1*, *aSMA* and *Postn* were assessed by RT-PCR. Data are shown as mean ± SEM and representative of three biological replicates. All reactions were performed in technical triplicates and normalized to GAPDH. Scale bar =100μm. *P < 0.05, **P <0.01 using t-test.

To extend our investigation to the human species, we used primary human cardiac fibroblasts (hCF) to define the function of PRMT1. TGFβ stimulation in hCF enhanced the expression of the myofibroblast protein αSMA and stress fiber formation in control siRNA-transfected fibroblasts, but not in PRMT1 siRNA-transfected hCF, similar to mouse fibroblasts (Fig 5K-L). As αSMA-rich stress fibers in myofibroblasts are known to be mechanically linked to the extracellular matrix and generate contractile force^13, 14^, we performed gel contraction assays to test whether PRMT1 regulates the contractile function of myofibroblasts. Upon TGFβ treatment, the collagen gel loaded with control siRNA-transfected fibroblasts exhibited significant contraction compared to PBS-treated group (Fig. 5M, 1^st^ row, right vs. left side). In contrast, the PRMT1-depleted fibroblasts showed a much weaker capacity to contract the collagen gel (Fig. 5M-N), indicating fewer functional myofibroblasts were formed. Furthermore, TGFβ-induced expression of myofibroblast signature genes *αSMA* and *Postn* was suppressed by PRMT1 depletion (Fig. 5O-Q). These results demonstrate that PRMT1-depleted fibroblasts fail to transition towards the myofibroblast fate after stimulation, suggesting a crucial role of PRMT1 in the transition of cardiac fibroblasts to myofibroblasts.

### PRMT1 depletion disrupts transcriptional and epigenetic regulation during human cardiac fibroblast transition to myofibroblasts

To elucidate the molecular mechanisms by which PRMT1 regulates myofibroblast formation, we analyzed transcriptional changes induced by TGFβ in control and PRMT1 siRNA-transfected hCF (Fig. 6A, 6B). In control siRNA-transfected fibroblasts, we compared TGFβ-treated versus PBS-treated groups and observed that TGFβ stimulation initiated the fate transition to myofibroblasts by upregulating a panel of myofibroblast signature genes (Fig. 6C-F, group 1: TGFb*Control siRNA). These genes are crucial for distinguishing myofibroblasts from quiescent fibroblasts at the molecular level and facilitate their contractile, migratory, and secretory functions. Among these, contractile genes include those traditionally associated with smooth muscle cells, such as *SMA* and *Tagln*, that enable the contractile function of myofibroblasts, and genes encoding focal adhesion proteins (vinculin, paxillin, integrin avb3, focal adhesion kinase, and actin), which were upregulated to promote directional migration to sites of cardiac injury (Fig. 6C-D). Myofibroblasts also demonstrated greater expression of proinflammatory cytokines, consistent with their secretory function that elevates the inflammatory status in the heart^1^ (Fig. 6E). Furthermore, myofibroblasts exhibited a metabolic switch, increasing aerobic glycolysis and oxidative metabolism^15, 16^ (Fig. 6F). In stark contrast, PRMT1 siRNA-transfected fibroblasts displayed marked inhibition in the induction of genes falling into all four categories (Fig. 6C-F). Pathway analysis further substantiated an inhibitory role for PRMT1, where genes associated with actin skeletal organization, muscle structure development, and metabolism were the top differentially expressed pathways when TGFβ-treated control hCFs were compared to TGFβ-treated PRMT1-depleted hCFs, indicating that PRMT1-depleted fibroblasts display molecular characteristics of the non-activated fibroblasts (Supplemental Fig. S4).

**Figure 6.**
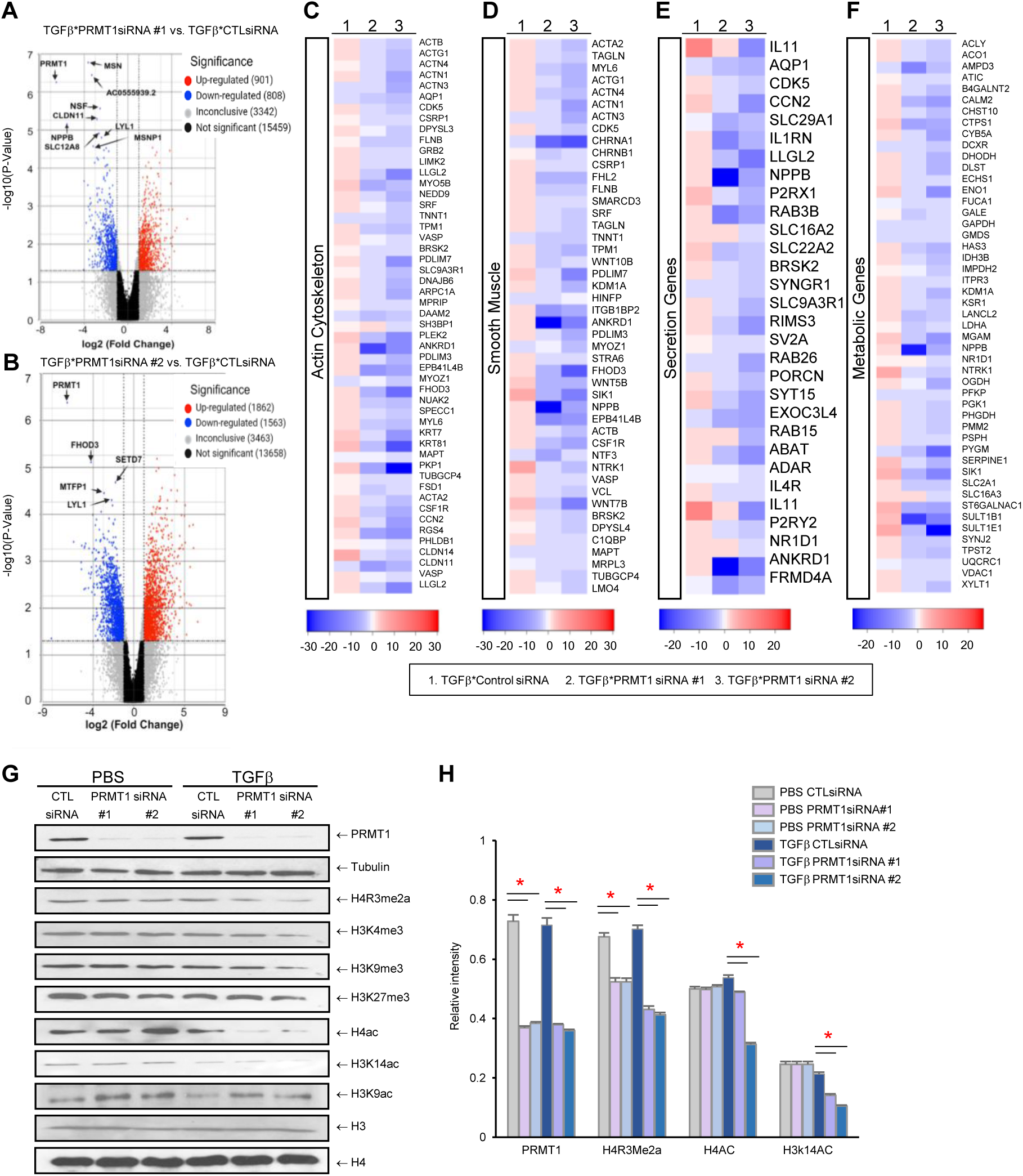
PRMT1 regulate transcriptional and epigenetic changes during fibroblast fate transition. **A-B.** Volcano plot showing PRMT1-regulated gene induction in hCF following TGFβ stimulated fibroblast to myofibroblast fate transition. TGFβ*Prmt1siRNA: TGFβ-treated and Prmt1 siRNA-transfected group compared to PBS-treated and Prmt1siRNA-transfected group. TGFβ*CTLsiRNA: TGFβ-treated and control siRNA-transfected group compared to PBS-treated and control siRNA-transfected group. **C-F.** Heatmaps of myofibroblast gene induction in hCF that was repressed by PRMT1 knockdown in actin cytoskeleton (C), smooth muscle (D), secretion (E) and metabolic genes (F). 1. TGFβ*Control siRNA, 2. TGFβ*Prmt1 siRNA #1, 3. TGFβ*Prmt1 siRNA #2. **G-H.** PRMT1 knockdown reduced H4R3me2a, H4Ac and H3K14 in myofibroblasts, revealed by immunoblotting. Quantification was conducted with Image J and shown in H. *P < 0.05 using t-test.

PRMT1 is also an epigenetic regulator that methylates histone H4R3, resulting in H4R3me2a, a histone mark associated with transcriptional activation^17^. To determine whether PRMT1 regulates the epigenetic status during fibroblast fate transition, we analyzed histone modifications in control and PRMT1 siRNA-transfected hCF (Fig. 6G-H). Depletion of PRMT1 diminished H4R3me2a in activated and non-activated hCFs (Fig. 6G, 3^rd^ row, 6H). Depletion of PRMT1 also reduced H4Ac and H3K14Ac, with no impact on H3K4me3, H3K9me3, or H3K27 (Fig. 6G-H), consistent with previous findings that H4R3me2a is required for H4 and H3K14 acetylation and the maintenance of an active chromatin^18^. Interestingly, PRMT1 depletion reduced H4Ac and H3K14Ac in activated fibroblasts, but did not alter their levels in the non-activated fibroblasts within the 48-72 hours period post-depletion.

These data demonstrate that PRMT1 exerts transcriptional and epigenetic regulations, and depletion of PRMT1 represses the activation of myofibroblast signature gene programs.

### PRMT1 binds to and methylates fibrillarin (FBL) in response to TGFβ stimulation

Besides transcriptional changes, we observed a profound change in ribosomal RNAs (rRNA), with a significant reduced mature 28s:18s ratio in PRMT1-depleted hCF (Fig. 7A-B). Importantly, the integrity of RNA samples was well-preserved, and the 18s levels remained stable across all groups. But the level of 28s was drastically decreased in PRMT1 siRNA-treated hCF after TGFβ stimulation (Fig. 7A-B). The 28s rRNAs are cleaved from the 47S precursor rRNA in the nucleolus, a membrane-less organelle specializes in rRNA processing^19^, suggesting a possibility that PRMT1 depletion disturbed rRNA processing during fibroblasts fate transition. To investigate the mechanistic link between PRMT1 and nucleolar function, we first examined the subcellular distribution of PRMT1. In primary hCF, PRMT1 mainly localized in the cytosol (Fig. 7C), which was distinct from a predominant nuclear localization in many epithelial and mesothelial cell types^20, 21^. Upon TGFβ stimulation, PRMT1 translocated into the nucleus (Fig. 7Ca), suggesting TGFβ-dependency of PRMT1 function in the nucleus and nucleolus in the activated fibroblasts.

**Figure 7.**
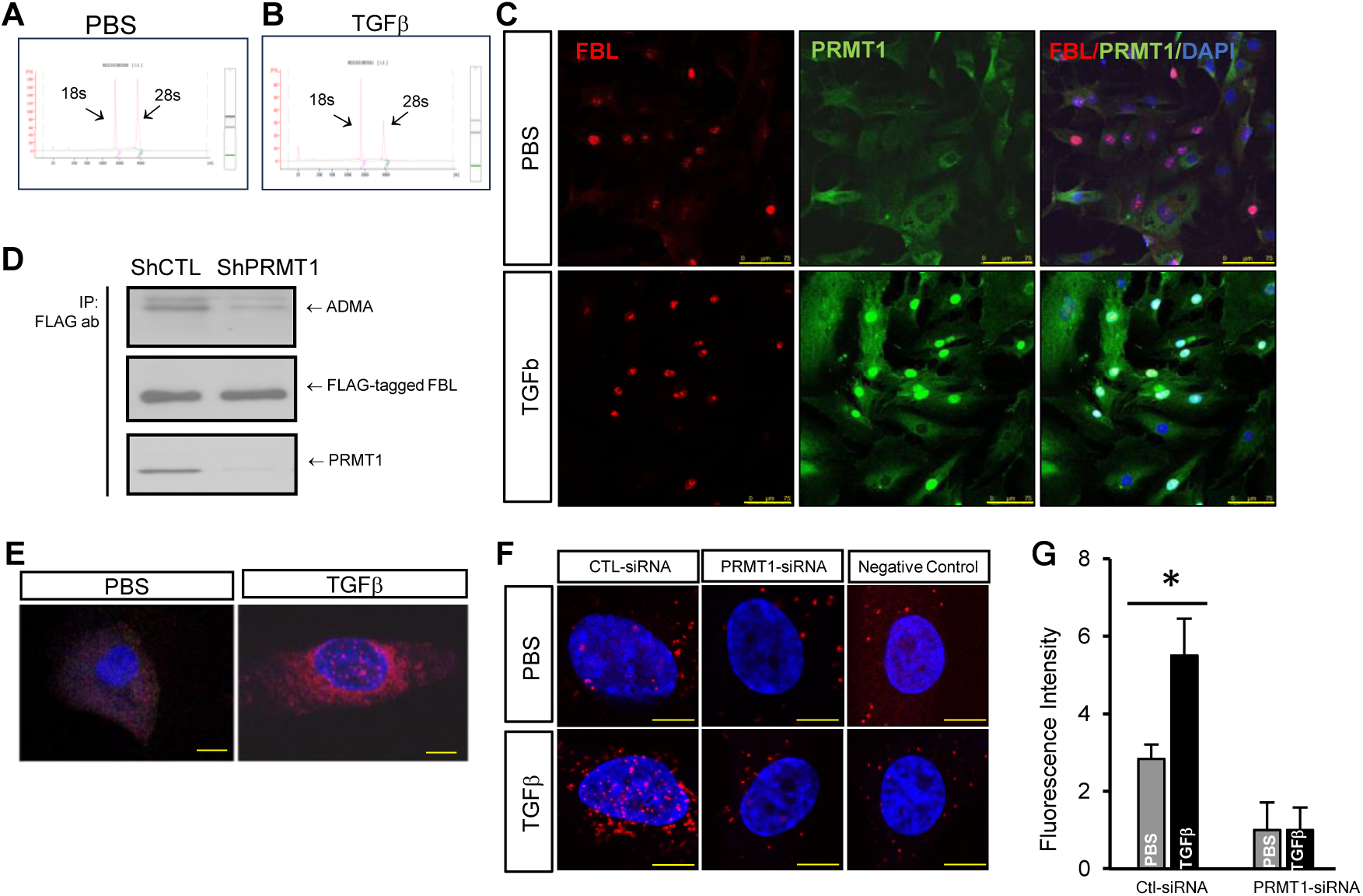
PRMT1 methylates FBL during cardiac fibroblasts to myofibroblasts transition. **A-B.** PRMT1 depletion caused drastic reduction of 28S rRNA. Representative bioanalyzer assay of total RNA samples from PBS or TGFβ-treated PRMT1-depleted hCF. **C.** PRMT1 translocates to the nucleus in response to TGFβ treatment in hCF. Immunohistostaining of PRMT1 and FBL in hCF treated with TGFβ or PBS. Scale bar = 75µm. **D.** Asymmetric dimethylarginine methylation (ADMA) of FBL reduced significantly in PRTM1 depleted cells. 293T stable cell lines expressing control shRNA or PRMT1 shRNA were transfected with vector expressing FLAG-tagged FBL, collected for immunoprecipitation (IP) using an anti-FLAG antibody, and analyzed by western blot (WB) using antibodies against asymmetric dimethylarginine methylation (ADMA), FBL or PRMT1. **E.** TGFβ stimulated PRMT1 interaction with FBL in hCF. The interaction between PRMT1 and FBL within the nucleus was detected by *in situ* proximity ligation assays (PLA) using PRMT1 and FBL antibodies and imaged using confocal microscope. DAPI nuclear counterstaining was used. Scale bar = 10 µm (×63 magnification). **F-G.** FBL arginine methylation was dramatically enhanced by TGFβ stimulation, and depended on PRMT1 in hCF. FBL methylation was detected by PLA assay with ADMA and FBL antibodies and imaged using confocal microscope. DAPI nuclear counterstaining was used. Negative control groups using FBL siRNA indicated that cytosolic punctate signal was non-specific. Scale bar=7.5 µm. Quantification of FBL methylation was conducted with puncta from five independent experiments and shown in G. *P< 0.05 using t-test.

Nucleolus consists of three compartments, the fibrillar component (FC), dense fibrillar component (DFC) and granular component (GC), separated by liquid-liquid phase separation (LLPS)^22^. The three compartments represent progressive stages of rRNA transcription, processing and ribosomal assembly. The initial steps of rRNA processing occur at DFC, which is established by nucleolar protein fibrillarin (FBL)^19, 23^. FBL contains a Glycine-Arginine-rich domain (GAR domain) in its N-terminal region that is known to be a substrate for methylation by PRMT1 in yeasts and in vitro reactions^24, 25^. However, it remains unclear whether FBL methylation occurs in response to physiological stimuli. Given that PRMT1 translocated to the nucleus upon TGFβ treatment, we hypothesized that TGFβ induces PRMT1 interaction with FBL and subsequent methylation. To investigate this, we first tested whether FBL methylation depends on PRMT1 in mammalian cells using cultured HEK 293T cells. Arginine methylation of FBL was detected using an antibody that recognizes asymmetrical dimethylated arginine residues (ADMA) catalyzed by type I PRMTs, including PRMT1 (Fig. 7D). When we depleted PRMT1 using shRNA, the arginine methylation level in FBL was dramatically decreased, indicating that PRMT1 is the major methyltransferase responsible for FBL arginine methylation in 293T cells (Fig. 7D). Next, we focused on hCF and investigated the spatiotemporal interaction between PRMT1 and FBL using proximity ligation assays (PLA) assay, which detects protein-protein interaction in situ. Initially, we found no detectable interaction between endogenous PRMT1 and FBL in control hCF (Fig. 7E, left panel). However, after TGFβ stimulation, a substantial association between PRMT1 and FBL was observed in the nucleus (Fig. 7E, right panel). We further examined the methylation of FBL in hCF using PLA assays with anti-FBL and arginine methylation antibodies. In control siRNA-transfected hCF, arginine methylation of FBL was detected in unstimulated fibroblasts, and TGFβ stimulation dramatically increased FBL methylation (Fig. 7F, left panels, top vs. bottom, 7G). However, this methylation signal was diminished in the PRMT1-depleted groups, suggesting that arginine methylation of FBL in hCF depends on PRMT1 (Fig. 7F, middle vs. left panels, 7G). The cytosolic punctate was unspecific signal, as illustrated in the negative control group, where FBL depletion ablated nuclear signals, but cytosolic signal remained unaffected (Fig. 7E, right panels). These findings indicate that FBL is methylated by PRMT1 in hCFs in response to TGFβ stimulation.

### PRMT1 regulates nucleolar morphology and activity during cell fate transition

Nucleoli dynamically respond to extracellular stimuli and environmental stress by undergoing shape and size changes, along with compositional alterations, to adapt to cell metabolism^19, 26, 27^. To determine whether nucleolar undergo changes during cardiac fibroblast to myofibroblast transition, we conducted an analysis of nucleolar dynamics during cardiac fibroblast fate transition. In untreated hCF, more than 70% of cells harbored irregular shaped nucleoli, as visualized by the nucleoli-specific proteins fibrillarin 1 (FBL) (Fig. 8A) and nucleolin (NCL) (Fig. 8B). Following TGFβ stimulation, a higher percentage of nucleoli acquired a spherical shape, shown in 32% of the cells (Fig. 8A-C). A small percentage of nucleoli exhibited a caps-like shape, suggestive of stress responses^28, 29^, and this subset was increased by TGFβ treatment (Fig. 8A-C).

**Figure 8.**
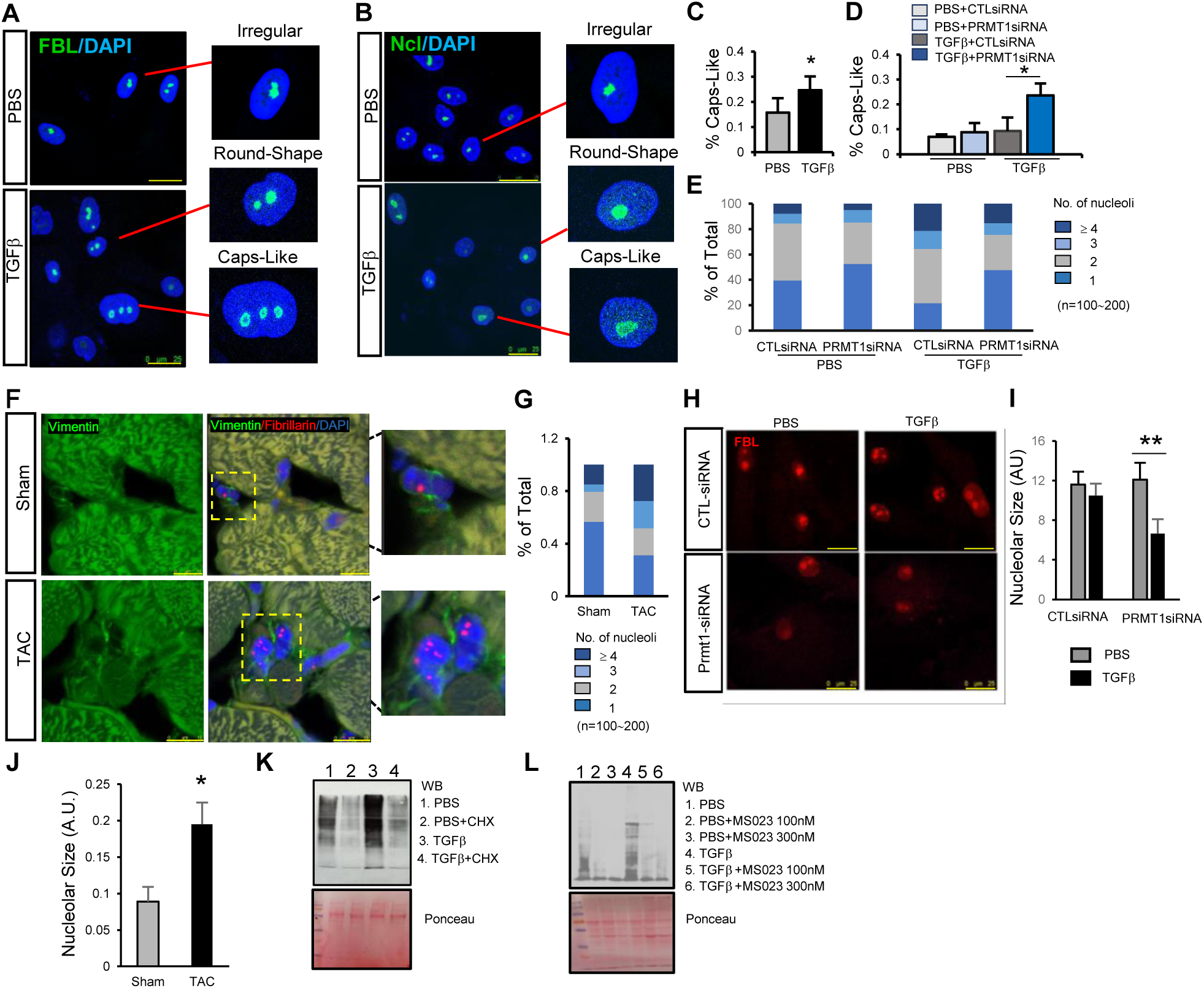
PRMT1 regulates nucleolar shape and size in cardiac fibroblasts. **A-C**. Cardiac fibroblast nucleolus display irregular or sphere shape, with a small percentage exhibiting caps-like structure (A-B). TGFβ treatment increased the percentage of nucleoli with a caps-like shape (C). Nucleoli shape was revealed by immunostaining using antibodies specific for nucleolar protein fibrillarin (**A**: FBL, green) and nucleolin (**B**: NCL, green) in hCFs after 48 hour of PBS or TGFβ treatment. DAPI counterstained the nucleus (Blue). Scale bar=25μm. The percentage of cap-like shape under TGFβ or PBS treatment was quantified in >100 cells per group and shown in C. **D.** PRMT1 knockdown increased the percentage of hCF with cap-like nucleoli. Nucleoli shape in control (CTL) siRNA or Prmt1 siRNA transfected hCF followed with TGFβ or PBS treatment was quantified in >100 cells per group. **E.** The number of nucleoli per nucleus in hCF increased following TGFβ-induced fate transition, but this increase was impaired by PRMT1 knockdown. hCFs were transfected with control (CTL) siRNA or Prmt1 siRNA followed with 48h TGFβ or PBS treatment. The number of nucleoli per cell was counted in 100-200 cells to generate the profile of nucleolar number in stacked bar for each group. **F-G.** The number of nucleoli per nucleus in cardiac fibroblasts increased in the mouse failing heart. Nucleoli in cardiac fibroblasts were revealed by immunostaining with anti-FBL antibody to label nucleolus (red) and anti-vimentin antibody to label fibroblasts (green) in the heart tissue sections from wild type mice receiving TAC (n=5) or Sham (n=5) operation. The number of nucleoli per cell was counted in 100-200 cells to generate the profile of nucleolar number in stacked bar and shown in G. **H-I.** PRMT1 depletion reduced nucleolar size in TGFβ-stimulated hCF. Immunostaining of FBL was performed in hCF with CTL siRNA or Prmt1 siRNA transfection followed with TGFβ or PBS treatment, and nucleolar size was quantified in >100 cells per group in I. **J.** Nucleolar size expanded in the mouse failing heart. The nucleolar size was quantified in >100 cardiac fibroblasts in each group using nucleoli revealed by anti-FBL (red) in vimentin-positive fibroblasts (green) in wild type mice receiving TAC (n=5) or Sham (n=5) operation. *P < 0.05, **P<0.01 from t-test. **K.** TGFβ treatment enhanced protein synthesis in hCF, as demonstrated by SUnSET assay using TGFβ or PBS treated hCF. Signal in both groups was inhibited by cycloheximide (CHX), an inhibitor of translation. **L.** MS023 reduced protein synthesis in PBS and TGFβ treated hCF, as demonstrated by SUnSET assay using TGFβ or PBS treated hCF with or without MS023.

LLPS fluidity imparts a dynamic, liquid-like behavior to the nucleolus, determining its shape and size^22^. PRMT1-catalyzed methylation of FBL occurs in the first 32 aa of its N-terminal GAR domain, which is crucial for self-association, phase separation and subsequent establishment of DFC ^30^. Methylation of arginine residue alters the hydrogen bonding property, thus exert critical control over protein-protein interaction and self-association^31^. To determine whether PRMT1-regulated methylation alters nucleolar dynamics and function, we analyzed nucleolar dynamics during cardiac fibroblast fate transition in control and PRMT1 siRNA-transfected cells. PRMT1 depletion impaired the maintenance of irregular shape in nucleoli, with approximately 44% of untreated cells showing irregular shapes as opposed to >70% in the control group (Fig. 8D). The caps-like, “stressed” nucleoli also increased significantly in PRMT1-depleted hCFs (Fig. 8D). Additionally, we observed changes in the number and size of nucleoli during the cardiac fibroblast transition to myofibroblasts. The majority of untreated hCF harbor one nucleolus (∼40%) or two nucleoli (∼40%) per nucleus (Fig. 8E). However, upon TGFβ stimulation, a higher percentage of cells hosted more than 4 nucleoli per nucleus, while only 20% retained one nucleolus per nucleus (Fig. 8E). PRMT1 depletion disrupted TGFβ-stimulated alteration and retained nucleoli number to a state similar to untreated hCFs (Fig. 8E). Interestingly, in mouse heart samples, nucleolar number in mouse cardiac fibroblasts demonstrated similar changes in TAC-induced heart failure samples when compared to sham-operated control, with the majority of sham cardiac fibroblasts harboring one nucleolus (>50%) or two nucleoli (∼25%) per nucleus, while TAC cardiac fibroblasts hosting more than 4 nucleoli per nucleus in a higher percentage (Fig. 8F-G). We further analyzed nucleolar size during fibroblast fate transition in cultured hCF. TGFβ stimulation did not induce overt changes in nucleolar size, but PRMT1 depletion decreased nucleolar size (Fig. 8H, 8I). In vivo, the nucleolar size of activated fibroblasts within TAC heart increased dramatically when compared with that in the sham group (Fig. 8J), potentially illustrating a chronic outcome undetectable in the cultured system.

The main function of nucleolus is ribosomal biogenesis and the assembly of protein synthesis apparatus, which prompted us to assess protein synthesis using SUnSET analysis, which detects puromycin-bound peptides released from the ribosome as the indicator of protein synthesis^32^. Cardiac fibroblast-to-myofibroblast transition induced by TGFβ was accompanied by an increased rate of protein synthesis (Fig. 8K, 3^rd^ vs. 1^st^ column). To investigate whether PRMT1 depletion in hCF affected new protein synthesis, we inhibited PRMT1 using MS023 during TGFβ-induced cell fate transition and demonstrated that the use of MS023 resulted in a reduction in new protein synthesis during the fate transition process (Fig. 8L, 2^nd^ & 3^rd^ vs. 1^st^ column, 5^th^ & 6^th^ vs. 4^th^ column). These findings demonstrate that PRMT1 depletion in cardiac fibroblasts impairs protein synthesis. Altogether, these data suggest that PRMT1 depletion impairs the nucleolar response to TGFβ-induced fibroblast fate transition structurally and functionally. The disruption in nucleolar dynamics and subsequent inhibition of protein synthesis may play critical roles in the impaired myofibroblast formation observed upon PRMT1 inhibition, shedding light on the importance of PRMT1 in the chronic fibrotic processes.

### FBL is essential for fibroblast to myofibroblast fate transition

FBL has been extensively studied for its role during mitosis in control of nucleolar disassembly and ribosomal biogenesis^33–36^. However, its involvement in fibroblast fate transition remains unknown. To elucidate the function of FBL in cardiac fibroblast transition to myofibroblasts, we depleted FBL in primary hCF using siRNA and stimulated fate transition with TGFβ (Fig.9A).

**Figure 9.**
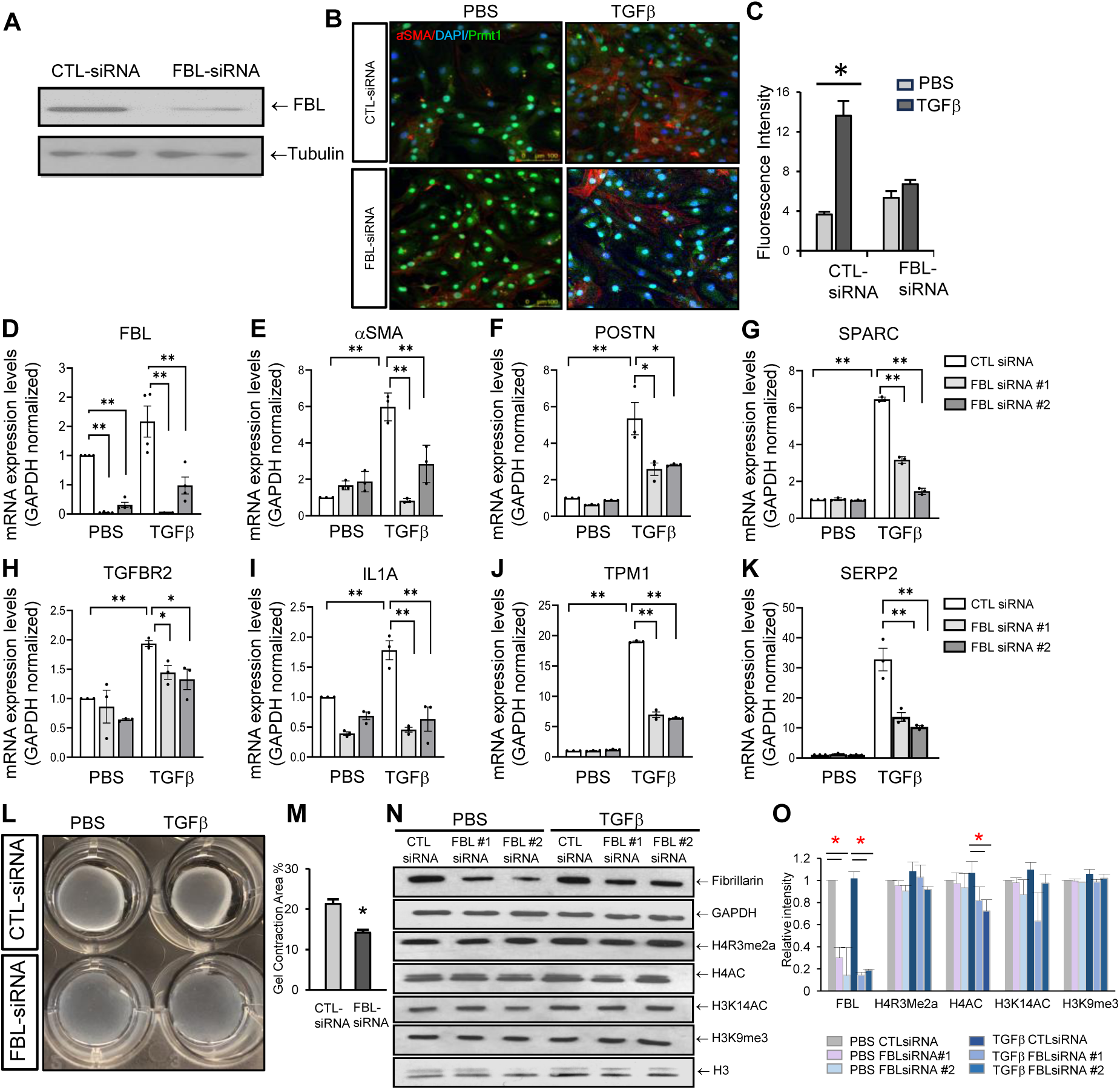
FBL is essential for fibroblast to myofibroblast fate transition. **A-C.** FBL depletion inhibited TGFβ-induced expression of myofibroblast marker αSMA in hCF. FBL is efficiently depleted in hCF by a pool of two independent siRNA (A). Western blot analysis of FBL in hCF transfected with control (CTL) or FBL siRNA and α-tubulin as loading control. Immunofluorescent signal of myofibroblast marker αSMA in FBL siRNA or control (CTL) siRNA-transfected hCF with TGFβ or PBS treatment was quantified and shown in C. Scale bar=50μm. **D-K.** The induction of myofibroblast marker genes *aSMA, Postn, Sparc, Tgfbr2, Il1a, Tpm1* and *Serp2* was repressed by FBL knockdown using two independent siRNAs in TGFβ-treated hCF. The mRNA levels of PRMT1 and myofibroblast marker genes were assessed by RT-PCR. Data are shown as mean ± SEM and representative of three biological replicates. All reactions were performed in technical triplicates and normalized to GAPDH. **L-M.** FBL depletion impaired contractile force generation in hCF. Collagen contractility assay of cultured hCF was conducted and the area of the gel was quantified and shown in M. **N-O**. FBL knockdown did not alter histone modifications in PBS or TGFβ-treated hCF. Quantification was conducted with Image J and shown in O. *P < 0.05, **P <0.01 using t-test.

Remarkably, FBL depletion significantly impaired the expression of myofibroblast protein αSMA and reduced stress fiber formation compared to control siRNA-transfected hCF after TGFβ stimulation (Fig. 9B-C). Subsequently, we assessed the transcriptional changes in FBL-depleted hCF during fibroblast-to-myofibroblast transition. FBL depletion repressed the transcriptional induction of myofibroblast signature genes involved in actin cytoskeleton organization, matrix formation and secretory function upon TGFβ stimulation (Fig. 9D-K), similar to changes observed in PRMT1-depleted hCF. Next, we determined whether FBL regulated the acquisition of contractile function, characteristic of myofibroblast activity. FBL depletion disrupted contractile force generation, as revealed by gel contraction assays using control and FBL-depleted hCF following TGFβ induction (Fig. 9L-M), in line with the transcriptional repression of actin cytoskeleton genes. We also wondered whether FBL depletion caused any epigenetic changes akin to PRMT1. There was apparent alteration in PRMT1-dependent H4R3 methylation in control or activated hCFs when FBL was depleted, but a moderate reduction in H4 acetylation was observed in the activated hCF (Fig. 9N-O). These discoveries highlight FBL as a new regulator of fibroblast fate transition to myofibroblast in hCFs and demonstrate that FBL is indispensable for the pathological induction of myofibroblast genes.

## DISCUSSION

In this study, we unveiled a previously unrecognized role of PRMT1 in promoting pathological cardiac fibrosis. Cardiac fibrosis profoundly contributes to cardiac dysfunction and heart failure, however, clinical interventions that effectively target cardiac fibroblasts and mitigate or reverse heart tissue fibrosis are still limited. Here we propose a new strategy that directly targets cardiac fibroblasts via blocking myofibroblast formation through inactivation of PRMT1. We demonstrated that genetic deletion of Prmt1 in activated cardiac fibroblasts almost abolished myofibroblast formation in the injured heart, which consequently inhibits cardiac fibrosis, reduces adverse remodeling, and preserves contractile function. PRMT1 is a druggable enzyme. Inspired by the tumor-promoting activity of PRMT1 in various cancer types, efforts have been made to develop small molecules that inhibit PRMT1 activity in vivo, such as MS023 and GSK3368715. Therefore, we investigated the potential of the PRMT1 inhibitor MS023 for treating cardiac fibrosis in TAC-induced mouse model of heart failure. A four-week treatment regimen with low dose significantly improved contractile function and reduced cardiac fibrosis in failing hearts at week 3 and 4. Another one-week treatment regimen with high dose MS023 also led to a notable mitigation of cardiac functional deterioration and reduction in cardiac fibrosis, with beneficial effects sustained for 3 weeks after drug withdraw. To the best of our knowledge, this study is the first to describe a protective role for PRMT1 inhibition in cardiac fibrotic remodeling. This study also present PRMT1 as a potential treatment target for cardiac fibrotic conditions.

Cardiac fibroblasts are the most abundant cell type in the heart, while cardiomyocytes constitute the largest cellular volume and generates the contractile force enabling cardiac output. The role of PRMT1 in cardiomyocytes have been determined previously using transgenic mouse models with constitutive *Prmt1* deletion, whereby loss of PRMT1 impaired cardiomyocyte maturation associated with defective splicing and deficient CaMKII methylation. However, cardiac functional assessment in these studies was also complicated by pre-existing myocyte maturation defects. Inspired by the beneficial effects of MS023 in TAC-induced heart failure mouse models, we deleted *Prmt1* in adult cardiomyocyte using an inducible *Myh6^MerCreMer^*, which revealed PRMT1 deletion as an anti-hypertrophy pathway (in a separate manuscript). Thus, MS023-induced anti-hypertrophic effects on cardiomyocytes observed in this study could be caused by either fibroblast-myocyte crosstalk or myocyte-intrinsic response. MS023-mediated inhibition of PRMT1 may act on multiple cell types in the heart to elicit beneficially effects.

The role of PRMT1 in cell fate transition have been described in multiple studies. PRMT1 promotes epithelial-to-mesenchymal transition (EMT) in breast and lung cancer cells through facilitating TGFβ signaling by Smad7 and Twist methylation^37–39^. PRMT1 also methylates EZH2 to regulate its repressor activity and methylates H4R3 to activate ZEB1 promoter^40, 41^. During heart development, PRMT1 promotes epicardial cell fate transition via controlling p53 turnover^20^. Here we demonstrate that PRMT1 promotes cardiac fibroblast to myofibroblast fate transition that involves FBL methylation. PRMT1 responds to TGFβ stimulation in cardiac fibroblasts and translocates to the nucleus, where it interacts with and methylates FBL. FBL acts as a mediator of the pro-fibrotic function since cardiac fibroblasts with FBL depletion were impaired in myofibroblast gene induction and contractile force generation. Considering the pleiotropic roles of PRMT1 in splicing, signaling, and transcription, PRMT1 may elicit FBL-independent pathways in cardiac fibroblasts fate transition to promote myofibroblast formation.

FBL is a key nucleolar component responsible for rRNA processing and pre-ribosomal assembly^42, 43^. Here we revealed new roles of FBL in fibroblast transition to myofibroblast in cardiac fibroblasts, where FBL depletion repressed the induction of myofibroblast genes responsible for actin cytoskeleton organization, leading to impaired contractile competence. Given its predominant localization in the nucleoli and regulatory roles in rRNA, the functional requisite for FBL in cell fate transition could arise from multiple tasks FBL performs. FBL-dependent ribosomal biogenesis is critical for protein synthesis, including the generation of transcriptional complex components that drive the myofibroblast transcription program, similar to the way that FBL regulates translation during neural stem cell differentiation^44^. FBL may also instruct the processing and assembly for cell type-specialized ribosome^45–47^, or modulate ribosomal transcription rate, which could affect cell fate determination^35, 48, 49^. These mechanisms, demonstrated in neural stem cells, cancer, ES or germ cells, await further investigation in the fibroblast fate transition process.

FBL localizes to the nucleolus, a phase-separated cell condensates in the nucleus that display dramatical changes in size, shape, and number in cancer, heart disease, and aging. Active and healthy nucleoli have irregular shapes and a fibrillar structure^28^. Under stress conditions, such as cytotoxic treatment, nucleoli become spherical, and nucleolar proteins form a ring delineating the edge of the sphere^26, 28, 50^. FBL is one of the nucleolar proteins that localized on the edge of the ring and visually manifests as a cap-like structure in stressed cells^28^. This nucleolar stress response was best characterized in cancer cells, where imaging quantification protocols were streamlined to facilitate anti-cancer drug screening^29, 51^. But nucleolus is also highly heterogenous among different cell and tissue types. Even in different cancer cell types, the size and number of nucleoli vary, ranging from one to six in documented cell types^52^. Knowledge of nucleolus in the heart are mainly collected from cardiac myocytes. The size and number of nucleoli in cardiomyocyte positively correlate with heart weight in human heart, and this increase in nucleoli size was thought to compensate for the increased demand for protein synthesis in a hyper-functional stressed heart^53, 54^. In the current study, we provide a morphological analysis of nucleolar structure in cardiac fibroblasts, where nucleolus respond to TGFβ stimulation from irregular shape to spheres, with 20-30% displaying caps-like arrangements that suggested nucleolar stress, suggesting that nucleoli undergo morphological changes during fibroblast transition to myofibroblast. Depletion of PRMT1 further increased the percentage of caps-like nucleoli, indicating augmentation of nucleolar stress that is consistent with loss of myofibroblasts after *Prmt1* deletion in vivo. Depletion of PRMT1 also reduced nucleoli size during TGFβ-stimulated fate transition, and inhibited protein synthesis. A change in nucleoli size in fibroblasts has been studies in aging. In healthy individuals, nucleoli size correlates with age in primary fibroblasts. Restricting nucleolar size can extend lifespan in multiple species^55, 56^, while nucleolar expansion in fibroblasts was documented in premature aging conditions. In diseased fibroblasts of mouse failing hearts, we observed a dramatic increase in nucleolar size, suggesting the expansion of nucleoli as a potential indicator of fibroblast pathology. Whether limiting nucleolar size would offer benefits for the heart is a question that requests future evaluation. A comprehensive assessment of nucleolar shape, number and size for cardiac fibroblast in healthy and diseased individuals would also be informative to evaluate whether nucleolar morphology may serve as a biomarker for the cardiac fibrotic process.

## DATA AVAILABILITY

All data generated or analyzed during this study are included in this published article and its Supplementary Information. RNA-sequencing data have been deposited at GEO and are publicly available as of the date of publication (Accession numbers: GSE255521).

## ACKNOWLEDGEMENT

This research was supported by American Heart Association Beginning Grant-in-Aid (16BGIA26540000, to J.X.), American Heart Association Postdoctoral Fellowship (to J.Q.), Rose Hills Foundation Research Fellowship (to J.X.), USC Stem Cell Center Audrey Streedain Regenerative Medicine Initiative Grant (to J.X. and R.S.) and Start-up fund from the Provost at USC (to J.X.). This work utilized the NMR Spectrometer Systems at Mount Sinai acquired with funding from the NIH’s SIG grants 1S10OD025132 and 1S10OD028504.

## AUTHOR CONTRIBUTIONS

J.Q. designed and conducted the experiments and wrote the manuscript. O.J.W. performed mouse cardiac fibroblast isolation and analysis. Y.C. and M.L. analyzed and interpreted RNAseq data. Y.S. and J.J. provided the MS023 for in vivo and in vitro studies. R.S. contributed to echocardiography data analysis. J.X. designed experiments, interpreted results, and revised the manuscript with contributions from all authors.

## COMPETING INTERESTS

J.J. is a cofounder and equity shareholder in Cullgen, Inc., a scientific cofounder and scientific advisory board member of Onsero Therapeutics, Inc., and a consultant for Cullgen, Inc., EpiCypher, Inc., Accent Therapeutics, Inc, and Tavotek Biotherapeutics, Inc. The Jin laboratory received research funds from Celgene Corporation, Levo Therapeutics, Inc., Cullgen, Inc. and Cullinan Oncology, Inc..

## Notes

### Competing Interest Statement

Jin Jian is a cofounder and equity shareholder in Cullgen, Inc., a scientific cofounder and scientific advisory board member of Onsero Therapeutics, Inc., and a consultant for Cullgen, Inc., EpiCypher, Inc., Accent Therapeutics, Inc, and Tavotek Biotherapeutics, Inc. The Jin laboratory received research funds from Celgene Corporation, Levo Therapeutics, Inc., Cullgen, Inc. and Cullinan Oncology, Inc..

