## Supplemental Materials for "Inactivation of PRMT1 inhibits cardiac fibrosis via transcriptional regulation and perturbation of FBL activity in fibroblast-to-myofibroblast transition"

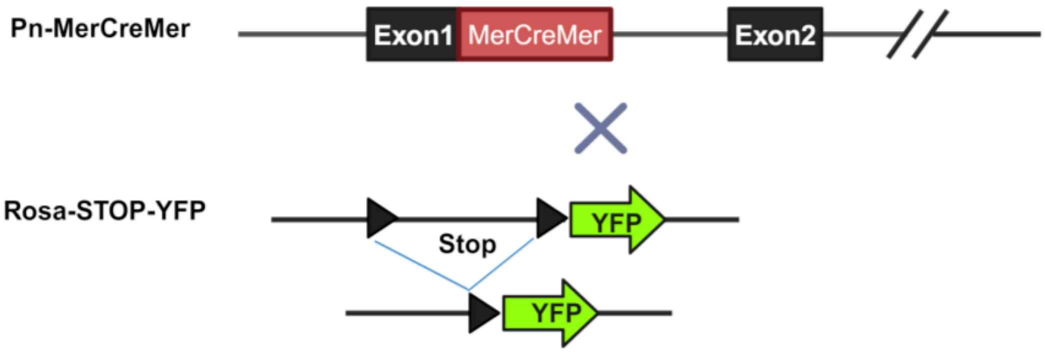


**Supplementary Figure S1. Generation of myofibroblasts lineage tracing mice.** Schematic representation of the Periostin (*Pn*) genetic locus with a tamoxifen-regulated MerCreMer (MCM) cDNA cassette inserted into exon 1 (E1). $Pn^{MCM}$ was crossed with Rosa26 reporter mice containing loxP sites flanking a stop cassette upstream of YFP ($ROSA^{STOP\text{-}YFP}$) to allow for Cre-dependent lineage tracing of myofibroblasts.

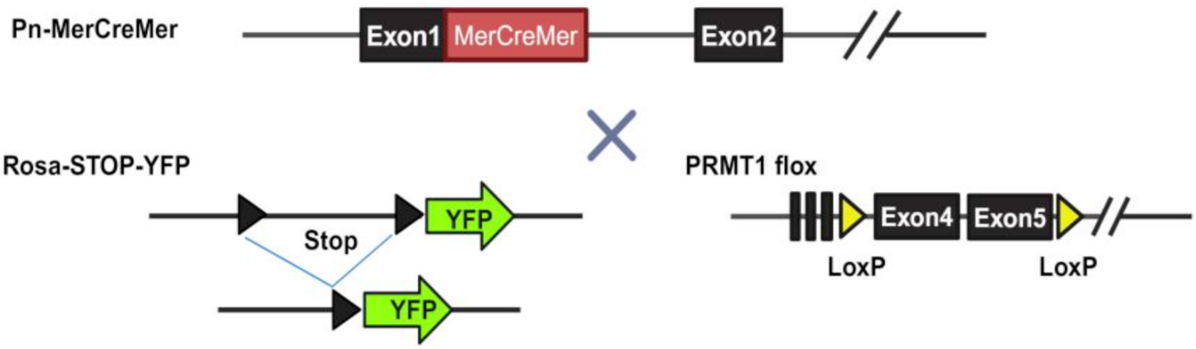


**Supplementary Figure S2. Generation of myofibroblasts-specific PRMT1 deletion mice.** Schematic representation of the tamoxifen-inducible $Pn^{MCM}$ crossed with $ROSA^{STOP\text{-}YFP}$ reporter mice and $Prmt1^{fl/fl}$ mice to generate myofibroblast-specific deletion of *Prmt1* and concurrent tracking of myofibroblasts by YFP.

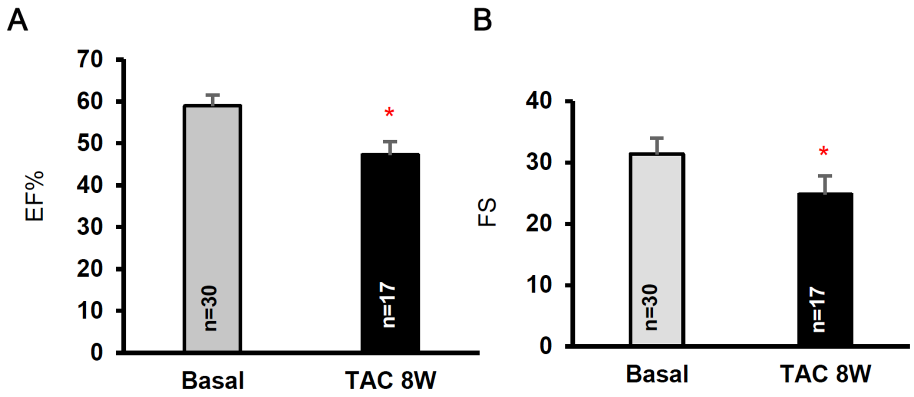


**Supplementary Figure S3. Induction of heart failure with TAC surgery in WT mice.** Echocardiographic assessment of left ventricular function of mice at baseline (n=30) and 8 weeks after TAC surgery (n=17). **A.** Measurement of ejection fractioning (EF%). **B.** Measurement of fractional shortening (FS). Results are reported as the mean ± SEM in each group. (*$P < 0.05$ from a t-test.)

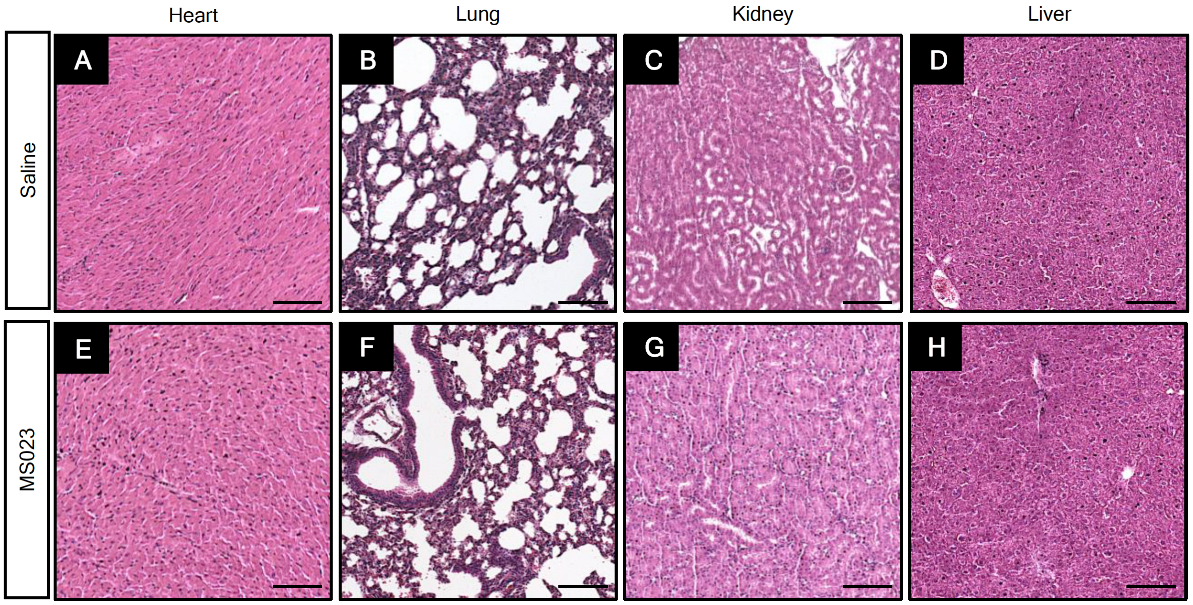


**Supplementary Figure S4. Histological analysis of major organs from mice receiving MS023 or saline treatments.** Hematoxylin and eosin (H&E) stained images of major organs from mice treated with MS023 and saline are used to assess local and systemic toxicity. The images of heart (A, E), lung (B, F), kidney (C, G) and liver (D, H) exhibit no overt morphological changes in MS023-treated groups. Scale bar=100µm.

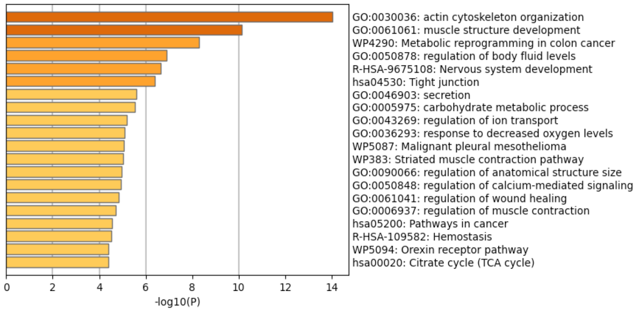


**Supplementary Figure S5**. Top 20 enriched Gene Ontology (GO) terms for differentially expressed genes when TGFβ-treated control siRNA transfected hCFs were compared to TGFβ-treated PRMT1 siRNA transfected hCFs.

**Supplementary Table S1. siRNA sequences**

| siRNA | Target Sequence |
|---|---|
| Prmt1siRNA #1 | CCC GTT CTG CCT GCA AGT GAA |
| Prmt1siRNA#2 | CAC CAT CGA CCT GGA CTT CAA |
| Fibrillarin siRNA #1 | ACA CTT TGT GAT TTC CAT TAA |
| Fibrillarin siRNA #2 | ATC GAG GAT GCT CGA CAC CCA |
| Control siRNA | AAT TCT CCG AAC GTG TCA CGT |

**Supplementary Table S2. Antibodies**

| Primary Antibody | Species | Dilution | Manufacture |
|---|---|---|---|
| PRMT1 | Rabbit | 1:100 IF | Cell signaling (#2449) |
| Asymmetric di-methyl arginine antibodies | Rabbit | 1:1000 WB | Cell signaling (#13522) |
| Fibrillarin | Rabbit | 1:1000 WB<br>1:50 IF | Cell signaling (#2639) |
| Fibrillarin | Mouse | 1:1000 WB<br>1:200 IF | Abcam (ab218846) |
| Fibrillarin | Rabbit | 1:50 IF | Thermo Fisher (A303-891A) |
| FLAG | Mouse | 1 µg FLAG antibody/500µl IP buffer | Millipore Sigma (B3111) |
| Nucleolin | Rabbit | 1:200 IF | Cell signaling (#14574) |
| α-Actin | Mouse | 1:50 IF | Millipore Sigma (#113200) |
| alpha Tubulin | Mouse | 1:1000 WB | Santa Cruz (#sc-8035) |
| Vimentin | Mouse | 1:200 IF | BD Biosciences (#550513) |
| CD31 | rat | 1:200 IF | BD Biosciences (#550274) |
| GFP | chicken | 1:200 IF | Abcam (#ab13970) |
| Secondary Antibody | Species | Dilution | Manufacture |
| HRP-conjugated goat anti-rabbit IgG (H+L) | Goat | 1:5000 WB | Jackson Immuno Research Laboratories (#111-035-003) |
| HRP-conjugated goat anti-mouse IgG (H+L) | Goat | 1:5000 WB | Jackson Immuno Research Laboratories (#115-035-003) |
| Goat anti-mouse IgG Alexa Fluor™ 594 | Goat | 1:200 IF | Thermo Fisher (A-11005) |
| Goat anti-rabbit IgG Alexa Fluor™ 594 | Goat | 1:200 IF | Thermo Fisher (A-11012) |
| Goat anti-mouse IgG Alexa Fluor™ 488 | Goat | 1:200 IF | Thermo Fisher (A-11001) |
| Goat anti-rabbit IgG Alexa Fluor™ 488 | Goat | 1:200 IF | Thermo Fisher (A-11008) |

**Supplementary Table S3. RT-PCR Primers**

| Gene | Species | Sequence 5' - 3' |
|---|---|---|
| IL1A-Forward | Human | TGTATGTGACTGCCCAAGATGAAG |
| IL1A-Reverse | Human | AGAGGAGGTTGGTCTCACTACC |
| TGFBR2-Forward | Human | GTCTGTGGATGACCTGGCTAAC |
| TGFBR2-Reverse | Human | GACATCGGTCTGCTTGAAGGAC |
| SPARC-Forward | Human | TGCCTGATGAGACAGAGGTGGT |
| SPARC-Reverse | Human | CTTCGGTTTCCTCTGCACCATC |
| POSTN-Forward | Human | CAGCAAACCACCTTCACGGATC |
| POSTN-Reverse | Human | TTAAGGAGGCGCTGAACCATGC |
| GAPDH-Forward | Human | GTCTCCTCTGACTTCAACAGCG |
| GAPDH-Reverse | Human | ACCACCCTGTTGCTGTAGCCAA |
| PRMT1-Forward | Human | CTTTGACTCCTACGCACACTT |
| PRMT1-Reverse | Human | GTGCCGGTTATGAAACATGGA |
| FBL-Forward | Human | GTCGAGGCGGAGGCTTTAG |
| FBL-Revers | Human | CATTCTTCCCCGACTGGTTTC |
| SERP2-Forward | Human | AGACTGGTCCTCGTCAACG |
| SERP2-Reverse | Human | CAGCATTGGCACTTGATAGGATT |
| TPM1-Forward | Human | TGGAGAGTCGAGCCCAAAAAG |
| TPM1-Reverse | Human | CATATTTGCGGTCGGCATCTT |
| Acta2 (aSMA)-Forward | Mouse | GTCCCAGACATCAGGGAGTAA |
| Acta2 (aSMA)-Reverse | Mouse | TCGGATACTTCAGCGTCAGGA |
| PRMT1-Forward | Mouse | GGGAGCCACTACCACTCAG |
| PRMT1-Reverse | Mouse | GTACGTGTATGACCCTTTTCCTT |
| Col1a1-Forward | Mouse | GCTCCTCTTAGGGGCCACT |
| Col1a1-Reverse | Mouse | CCACGTCTCACCATTGGGG |
| Col1a2-Forward | Mouse | TTCTGTGGGTCCTGCTGGGAAA |
| Col1a2-Reverse | Mouse | TTGTCACCTCGGATGCCTTGAG |
| Fn-EDA-Forward | Mouse | GTAACCAACATTGAT |
| Fn-EDA-Reverse | Mouse | GGCTCGAGTAGGTCAC |
| FN1-Forward | Mouse | CCCTATCTCTGATACCGTTGTCC |
| FN1-Reverse | Mouse | TGCCGCAACTACTGTGATTCGG |
| POSTN-Forward | Mouse | CAGCAAACCACTTTCACCGACC |
| POSTN-Reverse | Mouse | AGAAGGCGTTGGTCCATGCTCA |
| SPARC-Forward | Mouse | CACCTGGACTACATCGGACCAT |
| SPARC-Reverse | Mouse | CTGCTTCTCAGTGAGGAGGTTG |
| GAPDH-Forward | Mouse | ACGACCCCTTCATTGACCTG |
| GAPDH-Reverse | Mouse | AAATTTTCCTGGCTCCTGCT |

IL1A, Interleukin 1 Alpha; TGFBR2, Transforming Growth Factor Beta Receptor 2; SPARC, Secreted Protein Acidic and Cysteine Rich; POSTN, Periostin; GAPDH, Glyceraldehyde-3-

Phosphate Dehydrogenase; PRMT1, Protein Arginine Methyltransferase 1; FBL, fibrillarin; SERP2, Stress Associated Endoplasmic Reticulum Protein Family Member 2; TPM1, Tropomyosin 1; Col1a1, Collagen Type I Alpha 1 Chain; Col1a2, Collagen Type I Alpha 2 Chain; Fn-EDA, Fibronectin Extra Domain A ; FN1, fibronectin 1.

### METHODS

#### Animal sample size and randomization

The R26-stop-EYFP mutant mice (RRID:IMSR_JAX:006148) were purchased from the Jackson Laboratory, which have a loxP-flanked STOP sequence followed by the Enhanced Yellow Fluorescent Protein gene (YFP) inserted into the Gt(ROSA)26Sor locus. *Prmt1*$^{flox/flox}$ mice were provided by Dr. Stéphane Richard at McGill University. *Periostin*$^{MerCreMer}$ mice were provided by Dr. Jeff Molkentin at the University of Cincinnati. The number of animals used (n) was denoted in each test in the figures, including technical replicates when applicable. The mice were randomly selected from the cage and assigned to control and experimental subgroups as described in the text. The control and experimental groups were blinded to the operators of echocardiography and heart tissue analyses. The use of mice in our studies complied with the regulations of University of Southern California and National Institute of Health.

#### Isoproterenol osmotic pump implantation

Continuous infusion of isoproterenol with an osmotic pump was performed following the conventional procedure[57]. Briefly, mice were anesthetized with a mixture of inhaled isoflurane and oxygen-enriched air (1.25% during induction and 1% during maintenance). The surgical area, on the back of each animal between the shoulder blades, was cleaned and aseptically prepared. After a small incision, the skin was carefully separated from underlying connective tissues using blunt-ended scissors and an osmotic mini-pump (Alzet, model 1004) containing isoproterenol (Sigma Aldrich I6504) at 30 mg/kg per day dissolved in sterile 0.9% NaCl solution was implanted subcutaneously for delivery pharmacological agent for 14 days. Control group received the mini-pump that only contains sterile 0.9% NaCl solution. The incision was closed with 5–0 absorbable suture and surgical staples (Autoclip, Fine Science Tools). Buprenorphine SR (0.5 mg/kg) was subcutaneously administered 10 minutes before surgery and after 24 hours. Suture staples were removed 7 days after surgery.

**Transverse aortic constriction (TAC)**

TAC surgery was performed as previously described[58, 59]. In detail, male mice of 8-10 weeks of age were received endotracheal intubation first, which was connected to a Harvard volume-cycled rodent ventilator cycling at 125-150 breaths/minute and a tidal volume of 0.1-0.3ml. During the surgical procedure, anesthesia was maintained at 1.5-2% isoflurane with 0.5 - 1.0 L/min 100% O2. The aortic arch was exposed by opening the chest cavity at the level of the first intercostal space. The transverse aorta and a paralleling 27G needle were tied together with a 6–0 PROLENE polypropylene suture. The constriction of the aorta was produced after removal of the needle. The chest cavity was then closed with 6–0 silk suture. The skin was closed with 5–0 absorbable suture and Vetbond tissue adhesive (3M #1469SB). A sham group, undergoing surgery without aortic banding, was used as control.

**Echocardiography**

A Vevo 2100 high-resolution *in vivo* imaging system (VisualSonics Fujifilm) with a MS550S probe ‘high frame’ scan head was used for echocardiographic analysis. Mice were anesthetized with 1.0% isoflurane for M-mode imaging. Pressure gradients (60 to 90 mm Hg), an index of biomechanical stress, were determined by continuous wave Doppler on all animals that underwent TAC surgery. The left ventricular function was assessed by the M-mode scanning of the left ventricular chamber, standardized by two-dimensional, short-axis views of the left ventricle at the mid papillary muscle level.

**Cell culture**

Mouse cardiac fibroblasts were isolated from hearts of 10–14-week-old male mice. Ventricles of the hearts were cut into small pieces with scissors and digested with Liberase TH (#5401151001, Roche) in SADO-Mix Solution (20 mM HEPES-NaOH (pH 7.6), 130 mM NaCl, 3 mM KCl, 1 mM NaH2PO4, 4 mM Glucose, 1.5 mM MgSO4 in distilled water. The collected cells were cultured in DMEM with 10% FBS and 1% penicillin−streptomycin. Adult human cardiac fibroblasts were purchased from PromoCell (Cat#C-12375, Heidelberg, Germany) and cultured according to the manufacturer's instructions. Cardiac fibroblast fate transition to myofibroblast was stimulated with transforming growth factor-β1 (TGF-β1, #100-21 PeproTech, Cranbury, NJ, USA) at a final concentration of 20 ng/mL. Small interfering RNA (siRNA) transfection performed with the Lipofectamine RNAiMAX transfection reagent (#13778075, Invitrogen, Carlsbad, CA) according to the manufacturer's instructions. siRNAs were purchased from QIAGEN and listed in **Supplementary table S1.**

**Immunofluorescence staining**

As described previously[20] mouse hearts were washed in ice-cold PBS, fixed with 4% paraformaldehyde in phosphate-buffered saline (PBS), embedded in Tissue Tek (O.C.T. compound; Sakura), and frozen with dry ice. Serial sections (8-10 μm thick) of the tissue were prepared on a cryomicrotome at −21°C. After blocked with 5% goat serum in PBS +0.1% Triton X-100, tissue cryosections were incubated in primary antibodies overnight at 4°C. Antibodies and dilutions used were listed in the supplementary table 2. After three washes in PBS, sections were covered for 1 hour with the respective secondary antibodies (Alexa Fluor 488 or Alexa Fluor 594, Invitrogen). After washing in PBS, slides were mounted in Vectashield plus antifade medium with DAPI that counterstained the nucleus (Vector Laboratories). Images were taken under Leica confocal microscope and analyzed using ImageJ (RRID:SCR_003070). Mouse or human cardiac fibroblasts were cultured on the Millicell EZ slide (Millipore Sigma), then fixed and stained with the same protocol as the cryosection.

**Hematoxylin and eosin staining**

As described previously[60], mouse tissues were fixed with 4% paraformaldehyde in phosphate-buffered saline (PBS) and prepared into paraffin blocks, then sectioned into 8 μm thickness using a Leica microtome. Hematoxylin and eosin staining was performed for assessing tissue damage and observed under Leica microscope.

**Picrosirus Red staining**

The cryosections of mouse heart were stained with picrosirus red solution (0.1% direct red 80, #365548, Millipore Sigma) for 60 minutes and washed with two changes of saturated aqueous solution of picric acid (#P6744, Millipore Sigma). After quick dehydration, tissue sections were mounted and observed under polarized light microscopy (Leica).

**SUnSET assay**

SUnSET analysis[32] was used to assess protein synthesis in cultured human cardiac fibroblasts. Cells were incubated with 10 μg /mL puromycin (Sigma-Aldrich) for 15 min before or after incubation with 100 μg/mL cycloheximide for 10 min. All incubations were held at 37˚C. Cells were then pelleted and cell lysates analyzed via Western blot as described below. Mouse anti-puromycin monoclonal antibodies (Sigma-Aldrich) were used at a 1:2500 dilution. As a loading control, samples were stained with Ponceau S staining (0.5 % of Ponceau S dye and 1% of acetic acid, Sigma).

**SDS-PAGE, immunoblotting and immunoprecipitation.**

Proteins were extracted from cells using lysis buffer containing 50 mM Tris-HCl pH 7.5, 150 mM NaCl, 2mM EDTA, 0.1% NP-40, 10% glycerol and phosphatase/protease inhibitor cocktail. Protein concentration was measured using Bio-Rad Protein Assay (Bio-Rad Laboratories). Standard Western blot protocols, as previously published[20], were followed. For immunoprecipitation, cells lysates were pre-cleared with IgG and protein A/G Sepharose beads. Primary antibody was added and incubated overnight at 4°C. Immune complexes were precipitated with protein A/G Sepharose beads, washed three times with IP wash buffer (50 mM Tris-HCl pH 7.5, 150 mM NaCl, 2mM EDTA, 0.1% NP-40, 10% glycerol) and subjected to immunoblotting. Antibodies and dilutions were included in **Supplementary table S2.**

**Quantitative PCR**

Total RNA was isolated using a TRIzol-based (Invitrogen) RNA isolation protocol. RNA was quantified by Nanodrop (Agilent Technologies), and its quality were verified using an Agilent 2100 Bioanalyzer (Agilent Technologies). Samples required 260/280 ratios of more than 1.8, and sample RNA integrity numbers of more than 9 for inclusion. RNA was reverse transcribed using the Maxima H Minus cDNA synthesis kit (Thermo Scientific) according to the manufacturer's instructions. SYBR Green PCR Master Mix (Applied Biosystems, Foster City, CA) was used to amplify cDNA. GAPDH was used as loading control. Primers were listed in **Supplementary table S3.**

**RNA sequencing**

Total RNA extracted from human cardiac fibroblasts was enriched for poly(A) RNA and fragmented for library construction. RNA fragments were reverse transcribed followed by A-tailing and index adaptor ligation, then denatured and amplified on cBot. Library analysis and quality checks were performed on a LabChip GX. Sequencing was performed on an Illumina HiSeq 2000 platform with paired end reads of 100 bp. Analysis was performed using Partek Flow software (Partek Inc.). Differential expressed genes in each condition were identified using GSA. For gene ontology and pathway analysis, genes that were significantly up- or downregulated were compiled and analyzed using Ingenuity Pathway Analysis (QIAGEN Inc.).

**Proximity ligation assay**

To investigate the protein-protein interaction, an in situ Duolink® proximity ligation assay (PLA; Duolink Red starter kit for mouse; MilliporeSigma) was performed. Human cardiac fibroblasts were fixed in 4% paraformaldehyde. After blocking at 37 °C for 1 h, the cells were incubated with primary antibodies overnight at 4 °C. Primary antibody combinations were PRMT1 rabbit polyclonal and fibrillarin mouse monoclonal with 1:50 dilution. To detect protein arginine methylation[61], cells were incubated with primary antibodies against asymmetric di-methyl arginine motif (Cell signaling ) and fibrillarin (Abcam). Cells then were stained with anti-mouse MINUS probe (DUO92004) and anti-rabbit PLUS probe (DUO92002) that contains unique DNA strands. A ligase hybridization step followed by addition of polymerase for 100 minutes at 37 °C amplified the fluorescent probes then permit visualization of the red spots of proximity under Leica confocal fluorescence microscope. PLA signal was quantified as the total number of spots / nuclei. A total of 50 nuclei were counted for each group.

**Statistics**

Data are shown as the mean ± SEM. GraphPad Prism (version 10, GraphPad Software, La Jolla, California) was used for statistical analysis. One-tailed unpaired Student's t-test was used for 2-group comparison. We used one-way analysis of variance (ANOVA) followed by Tukey-Kramer's post-hoc test or two-way ANOVA followed by Bonferroni's post-hoc test for multiple comparisons. $P < 0.05$ was considered statistically significant.
